# Genomic structures and regulation patterns at HPV integration sites in cervical cancer

**DOI:** 10.1101/2023.11.04.564800

**Authors:** Vanessa L. Porter, Kieran O’Neill, Signe MacLennan, Richard D. Corbett, Michelle Ng, Luka Culibrk, Zeid Hamadeh, Marissa Iden, Rachel Schmidt, Shirng-Wern Tsaih, Glenn Chang, Jeremy Fan, Ka Ming Nip, Vahid Akbari, Simon K. Chan, James Hopkins, Richard A. Moore, Eric Chuah, Karen L. Mungall, Andrew J. Mungall, Inanc Birol, Steven J. M. Jones, Janet S. Rader, Marco A. Marra

## Abstract

Human papillomavirus (HPV) integration has been implicated in transforming HPV infection into cancer, but its genomic consequences have been difficult to study using short-read technologies. To resolve the dysregulation associated with HPV integration, we performed long-read sequencing on 63 cervical cancer genomes. We identified six categories of integration events based on HPV-human genomic structures. Of all HPV integrants, defined as two HPV-human breakpoints bridged by an HPV sequence, 24% contained variable copies of HPV between the breakpoints, a phenomenon we termed heterologous integration. Analysis of DNA methylation within and in proximity to the HPV genome at individual integration events revealed relationships between methylation status of the integrant and its orientation and structure. Dysregulation of the human epigenome and neighboring gene expression in *cis* with the HPV-integrated allele was observed over megabase-ranges of the genome. By elucidating the structural, epigenetic, and allele-specific impacts of HPV integration, we provide insight into the role of integrated HPV in cervical cancer.

## Introduction

Human papillomavirus (HPV), an 8-kilobase (kb), double-stranded, circular DNA virus, drives nearly all cervical cancers and a subset of head and neck cancers and anogenital cancers^1^. Vaccination against high-risk HPV types has effectively reduced cervical cancer rates^2^. However, cervical cancer incidence remains high in countries lacking vaccine availability^3^. During HPV-driven oncogenesis, the viral DNA commonly becomes integrated into the host cell genome^4^. HPV integration can lead to changes in the sequence content of HPV, viral gene copy number, and epigenetic regulation of the viral genome^5–8^. Genomic and epigenomic changes can also occur in adjacent human genomic regions^9–12^, and may have cancer-promoting consequences, as they often include oncogenes within or near the site of HPV integration^13^.

HPV integration is not a normal part of the HPV life cycle and occurs when double-stranded breaks in non-contiguous regions of the human genome are bridged using surrogate double-stranded DNA from the virus^8^. Integration occurs more frequently at regions with HPV microhomology, at fragile sites, or at sites that are actively transcribed^14^. HPV integration is also commonly found in regions dense with structural alterations^8,15^. Furthermore, the resolution of an unstable HPV-human concatemer can result in intermediate “onion-skin” structures with multiple replication bubbles that cause several human-HPV breakpoints across a genomic locus, which are referred to as an integration event^15^. The genomic consequences of HPV integration have thus far been difficult to study due to the limitations of short reads in capturing the complexity of human-HPV structures. However, recent advancements in long-read DNA sequencing technology have enhanced our ability to interpret repeating sequences, structural alterations, and DNA methylation signals^16^. Combined with enhanced haplotype phasing capabilities, long-read sequencing can therefore be used to resolve complex HPV-integrated genomes and methylomes^17^.

To investigate the structural changes resulting from HPV integration and their impacts on human and viral epigenetic regulation, we used long-read sequencing to characterize the genomes of 63 HPV-positive cervical cancer tumors: 43 from the HIV Tumor Molecular Characterization Project (HTMCP) and 20 from The Cancer Genome Atlas (TCGA). We found 504 integration breakpoints mapping to 130 integration events across 57 HPV-integrated cervical cancer samples. We grouped structural events associated with HPV integration into six categories: deletion, duplication, extrachromosomal circular DNA (eccDNA), translocation, repeat region, and multi-breakpoint integrations. Multi-breakpoint integrations were the most prevalent and contained complex genome rearrangements spanning up to four chromosomes and clustered integration breakpoints on loci spanning up to 8 megabases (Mb). Between breakpoints, we observed heterologous copies of the HPV genome, and variable HPV copy numbers. In the methylome, we observed frequent methylation of human genomic regions 5’ of the HPV integrants, whereas 3’ regions were unmethylated. HPV genes were also consistently hypermethylated in samples with integrated HPV compared to samples containing only episomal HPV. Differential methylation analysis of haplotypes identified large genomic regions in which HPV integration overlapped with haplotype-specific changes in methylation, some of which were also associated with allele-specific gene expression changes. Our analysis thus revealed previously unappreciated genomic and epigenomic dysregulation of human and viral genomes associated with HPV integration in cervical tumors.

## Results

### ONT whole genome sequencing

The HTMCP samples yielded an average of 106 gigabase pairs (Gb) of data (ranging from 58 - 153 Gb; median of 32X redundant coverage) and the TCGA samples yielded an average of 92 Gb (ranging from 31 - 186 Gb; median of 29X redundant coverage; Extended Data Fig. 1). Reads were longer in the HTMCP samples (median N50 of 19.6 kb; ranging from 14.3 - 34.1 kb) than the TCGA samples (median N50 of 12.8 kb; ranging from 0.786 - 27.2 kb; Extended Data Fig. 1). The proportion of reads from artifactual chimeric DNA fusions were determined using a spike-in lambda genome control and detecting human-lambda chimeras. As expected from the longer read lengths, the HTMCP samples had lower chimeric rates than the TCGA samples, and lower base error rates, indicative of higher quality reads (Extended Data Fig. 1). These differences in quality between cohorts may be due to the differences in the age of the tumor samples and the number of freeze / thaw cycles they had been subjected to.

### Long-read sequencing detection of HPV integration

The molecular and clinical properties of the 63 samples are shown in Fig. 1a (Supplementary Table 1). We observed that HPV integration events involved at least two double-stranded human genome breaks and two double-stranded HPV genome breaks (Fig. 1b). Breakpoints were grouped into an “integration event” if either of the following two conditions was met: (1) the HPV breakpoints co-occurred on one or more of the same reads, or (2) the breakpoints mapped within 500 kb of each other^18,19^ (see Methods). Applying these conditions, we detected 504 integration breakpoints belonging to 130 integration events across 57 samples with HPV integration (Fig. 1a; Supplementary Table 2). In six samples with deep and evenly distributed read coverage across the HPV genome, we did not detect HPV integration breakpoints, indicative of highly amplified episomal HPV.

**Figure 1:**
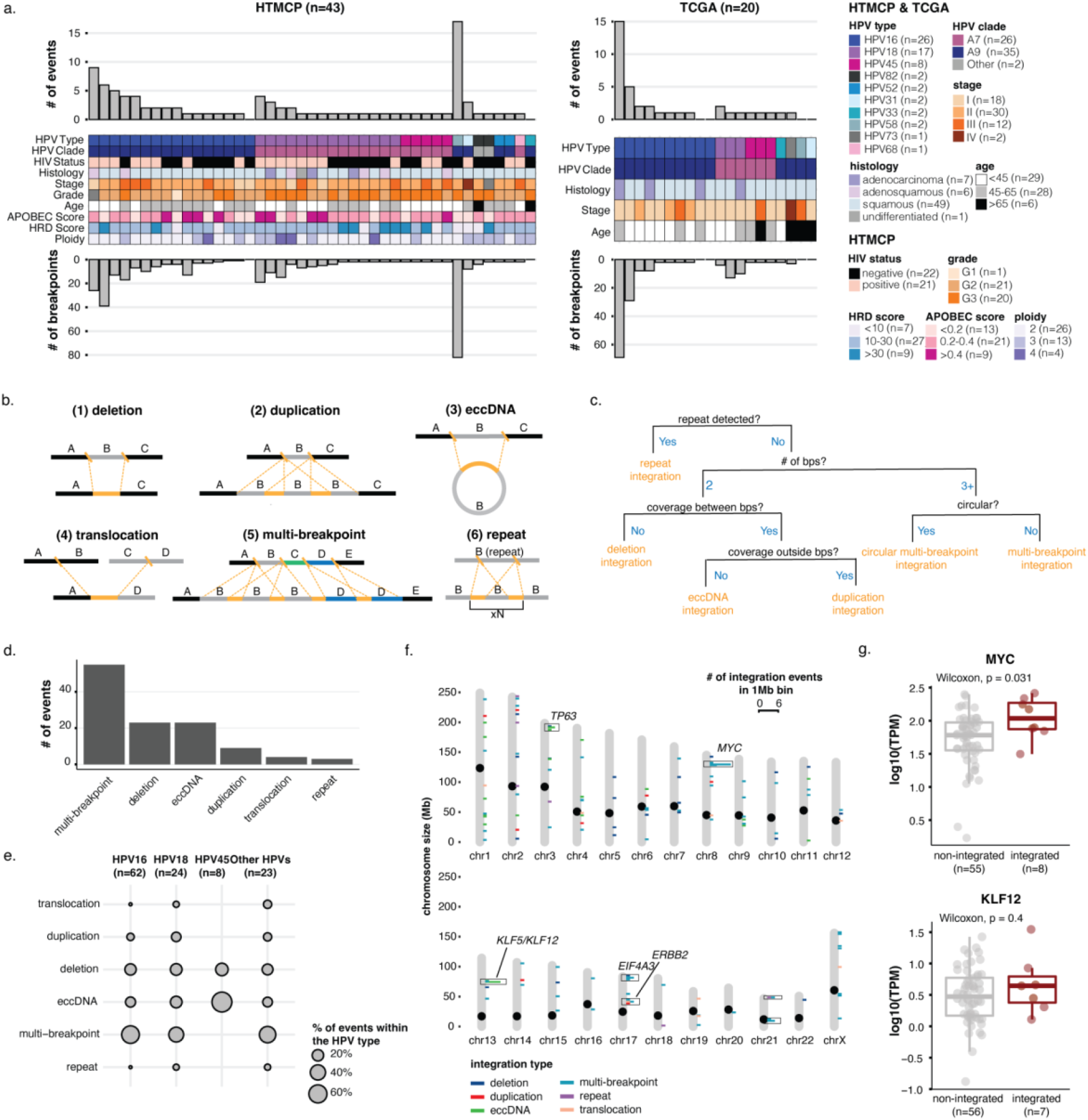
Detection and categorization of HPV integration events in cervical cancers. **(a)** The number of HPV integration breakpoints and events across the HTMCP and TCGA samples, and clinical and molecular characteristics of the patient samples. **(b)** Schematic illustrations of integration event categories. Orange segments indicate the insertion of HPV integrants. **(c)** Decision chart for categorizing HPV integration events into the listed structures (“bps” = “breakpoints). **(d)** The frequency of the HPV integration types across the samples. **(e)** The percentage of events belonging to each integration category for HPV16, HPV18, HPV45, and all other HPV types. **(f)** The genomic locations of the integration events across the cohort, coloured by integration type. Bins with 2+ integration events are indicated by boxes. Notable cancer genes within bins recurrently affected by integration are indicated. **(g)** Gene expression differences between samples with HPV integration within 1 Mb of *MYC* (top) and *KLF12* (bottom) compared to the remaining samples in the dataset. Box plots represent the median and upper and lower quartiles of the distribution; whiskers represent the limits of the distribution (1.5 IQR below Q1 or 1.5 IQR above Q3). *P* values were calculated using the Wilcoxon rank-sum test.

Next, we compared the number and genomic locations of HPV integration breakpoints identified in our long-read sequencing data to previous short-read sequencing data^19^ (Methods). Across the 41 HTMCP samples with HPV integration, 319 and 343 integration breakpoints were called in the short-read and long-read sequencing datasets, respectively (Supplementary Table 2 and Supplementary Table 3). Integration breakpoint calls using short and long reads were identical for 17 samples. Fourteen samples contained more calls in the long-read data, and 10 contained more calls in the short-read data (Extended Data Fig. 2a-b). Notably, two samples had highly discordant integration breakpoint calls, with an HPV58-integrated sample having 59 more calls in the long-read data (82 vs 23) and an HPV16-integrated sample having 53 more calls in the short-read data (56 vs 3; Extended Data Fig. 2a-b). We confirmed overlap between integration breakpoint calls between these two samples; 21/23 (91%) short-read calls in the HPV58-integrated sample were also reported in the long-reads, and 2/3 (67%) long-read calls in the HPV16-integrated sample were also reported in the short-reads. In the HPV58-integrated sample, 67/105 (64%) integration breakpoints called in either the long-read or short-read data were predicted to overlap repetitive regions, and 30/59 (51%) integration breakpoints resided in repetitive regions in the HPV16-integrated sample, suggesting that repetitive sequences may have caused the discordant calls. The HPV16-integrated sample also had lower read support for each breakpoint call (33/56; 59% with ten or fewer reads) than the HPV58-integrated sample (18/82; 22% with ten or fewer reads), indicating that the short-read integration breakpoint calls were of lower confidence, particularly at repetitive regions. As expected, the number of integration breakpoint calls per sample was correlated between technologies (Extended Data Fig. 2c; Spearman’s rank correlation, *R*=0.72, *p*=2.3 x 10^−7^). Both technologies detected similar proportions of breakpoint calls within CpG islands and exonic, intronic/intergenic, and repetitive regions. (Extended Data Fig. 2d). Thus, long-reads detected more integration breakpoints per sample, and discordance between long- and short-read technologies could be due to low-confidence mapping of short reads in repetitive regions.

### The six major categories of HPV integration events

Analysis of the HPV-integrated sequence alignment patterns revealed six categories of integration structures in the 57 integrated samples. These were deletion, duplication, extrachromosomal circular DNA (eccDNA), translocation, multi-breakpoint, and repeat region integrations, which we categorized using a custom workflow (Fig. 1b-c; Methods). Multi-breakpoint integrations were the most prevalent, comprising 55/130 (42%) of the total integration events in our dataset (Fig. 1d; Supplementary Table 2). Next were deletions (23/130) and eccDNA integrations (23/130), while duplication (9/130), translocation (4/130), and repeat region (3/130) integrations were rare (Fig. 1d; Supplementary Table 2). The remainder of the events (13/130) were unmatched single breakpoints that we were unable to categorize (Supplementary Table 2). The distributions of these integration categories differed between HPV types. For example, HPV16 was more likely to be involved in multi-breakpoint events than HPV18, whereas duplication integrations were rarer (Fig. 1e). HPV45 integrations were most frequently involved in eccDNA structures, which made up 75% (6/8) of its integration events (Fig. 1e).

We observed HPV integration hotspots (>2 events within 1 Mb bins) in the genome, most of which were dominated by a single HPV integration category (Fig. 1f). The two most frequently integrated hotspots were a region near *MYC* (8q24) and an intergenic region between *KLF5* and *KLF12* (13q22). In the 8q24 hotspot, multi-breakpoint events were predominant, whereas eccDNA events were predominant at 13q22 (Fig. 1f; Extended Data Fig. 3a-b). Four eccDNA events within 13q22 overlapped each other and contained a region shared by all four samples (Extended Data Fig. 3a). We compared the expression of *MYC* and *KLF12* in samples with and without HPV integration at these hotspots in the HTMCP and TCGA cohorts (Fig. 1g; Supplementary Table 4). *MYC* expression was increased in the integrated samples, while *KLF12* expression was not affected (Wilcoxon, *p*=0.031, Fig. 1g).

### HPV copy number heterogeneity within integration events

We defined an HPV integrant as the uninterrupted segment of the integrated HPV genome situated between two breakpoints. Examples of possible integrant structures are shown in Fig. 2a. An HPV breakpoint can be unambiguously linked to another HPV breakpoint when the entirety of the integrant is contained on a single read, and we refer to each unique pairing as a breakpoint pair (Supplementary Table 5). Cases with read support for inconsistent integrant sizes between a single breakpoint pair were referred to as heterologous integrants (Fig. 2b). Heterologous integrants existed in 24% (61/253) of breakpoint pairs (Fig. 2c; Supplementary Table 5). The maximum number of unique integrant structures found between a breakpoint pair was 15, occurring within an HPV16 multi-breakpoint integration event (Fig. 2c). Fig. 2d and Fig. 2e summarize the differing sizes of HPV integrant structures originating from each breakpoint pair in two samples. Both samples harbored heterologous integrants in breakpoint pairs originating from multi-breakpoint integration events. The heterologous integrants had read lengths consistent with integrants that contained a partial copy of HPV (n) plus additional complete HPV copies (n+1, n+2, etc.), as seen in event three of HTMCP-03-06-02238 (Fig. 2e). HTMCP-03-06-02058 provides an atypical example, where a portion of the HPV18 genome is deleted and the heterologous structures contain different copy numbers of the remaining HPV segment (Fig. 2d). The same portion of HPV was deleted in all breakpoint pairs in this integration event. These two events provide examples of the periodicity of integrant lengths in heterologous integrants, which we hypothesize arise through unequal amplification of the HPV genome during the amplification of HPV-human concatemers, particularly within multi-breakpoint events.

**Figure 2:**
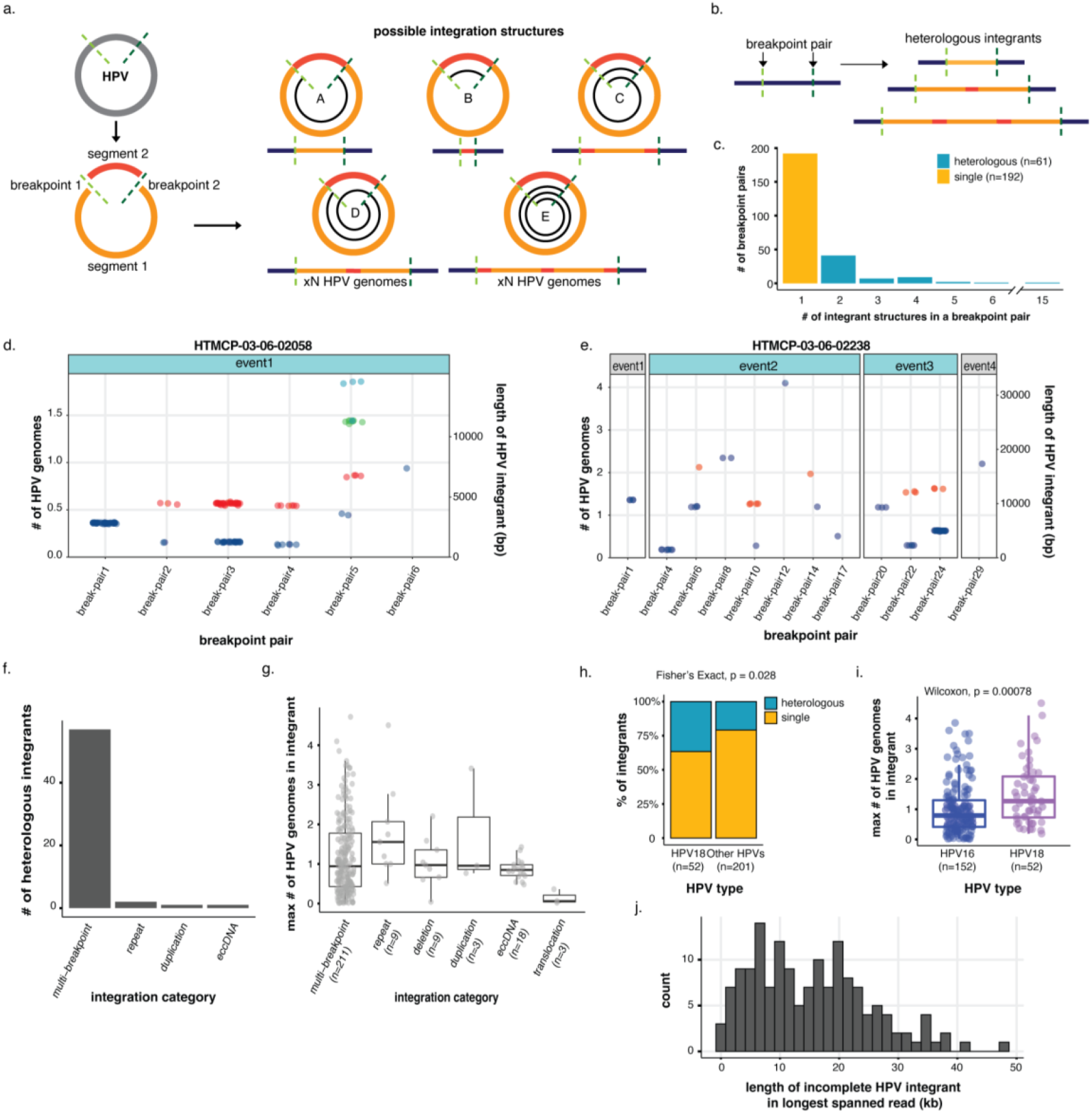
Heterologous structures of the HPV genome in single HPV integrants. **(a)** Schematic of possible structures within an HPV integrant. The spirals in structures A-E show the portion(s) of the HPV genome that could be contained within an integrant. **(b)** Schematic showing how several heterologous integrants can exist between a single breakpoint pair, with the size of the integrant varying by *n* HPV copies. The colors correspond to the regions of the HPV genome as depicted in (a). **(c)** The number of integrant structures between all identified breakpoint pairs within the cohort. Integrants with 2+ identified structures were classified as heterologous. **(d,e)** The sizes of the HPV integrants identified on individual reads in (d) HTMCP-03-06-02058 and (e) HTMCP-03-06-02238, grouped by breakpoint pairs. The multi-breakpoint events in these samples (colored blue) are the only ones with heterologous integrants. Individual points represent the size of the HPV integrant on individual reads, which are then grouped together by color if they do not differ in size by more than 300 bp. Each unique color in a breakpoint pair thus represents one unique integrant structure. **(f)** Frequency of heterologous integrants in the different integration categories. Categories that are not shown harbored no heterologous integrants. **(g)** The maximum number of HPV genomes found in an integrant across the different integration types. **(h)** The percentage of integrants from different HPV types that form single or heterologous structures. **(i)** The maximum size of the integrant structure in each breakpoint pair in HPV16 and HPV18 integrants, represented as the number of HPV genome copies. **(j)** Distribution showing the maximum number of HPV copies found in the longest-spanning read for each incomplete integrant. The x-axis shows the length of the longest-spanning read. Box plots represent the median and upper and lower quartiles of the distribution; whiskers represent the limits of the distribution (1.5 IQR below Q1 or 1.5 IQR above Q3). *P* values were calculated using the Wilcoxon rank-sum test and the Fisher exact test, as listed on the figure.

Breakpoint pairs from multi-breakpoint events made up 93% (57/61) of the heterologous integrants (Fig. 2f; Supplementary Table 5). Notably, heterologous integrants were not detected in deletion or translocation integration events, suggesting amplified copy number is associated with their formation (Fig. 2f). Translocation integration events involved smaller HPV integrants than the other integration categories, although only three complete translocation integrants were available (Fig. 2g). When comparing HPV types, HPV18 integrants were more frequently heterologous than other HPV types (37% vs. 21%; Fisher’s exact test, *p*=0.028; Fig. 2h) and contained significantly more HPV copies per integrant than HPV16 (Wilcoxon, *p*=0.00078; Fig. 2i).

We were unable to determine the complete length of the integrant for 153 breakpoints because no reads linked the detected breakpoint to another and thus they could not be paired. For each of these incomplete integrants, we identified the read that achieved the greatest span and used it to determine the minimum size of the integrant (Fig. 2j; Supplementary Table 5). Incomplete integrants often contained at least two HPV copies (mean of 1.9 HPV copies) and approximately half belonged to HPV16 events (77/153 breakpoints, 50%). The longest incomplete HPV integrant belonged to an HPV16 deletion event, which was supported by a read that spanned 48 kb (Fig. 2j). In contrast, the largest complete integrant between a breakpoint pair was 37 kb. Thus, even with long-read sequencing, we did not achieve complete resolution of HPV integrant lengths in 153/504 (30%) of HPV breakpoints.

### The three possible resolutions of two-breakpoint HPV integration events

Two-breakpoint integration events involving one chromosome, comprising 55/130 (42%) of our cohort, occur when one HPV integrant is inserted between two breaks in the human genome. These were associated with three integration types: deletions, duplications, and eccDNAs (Fig. 1b), which could be differentiated based on the alignment patterns of the HPV-containing chimeric reads. Integration-mediated deletions were characterized by a copy number loss between the integration breakpoints, in some cases accompanied by amplification of the region flanking the integration (Fig. 3a). In duplication and eccDNA integrations, the region between the breakpoints was amplified, but with differing patterns of read alignment outside the breakpoints (Fig. 3a).

**Figure 3:**
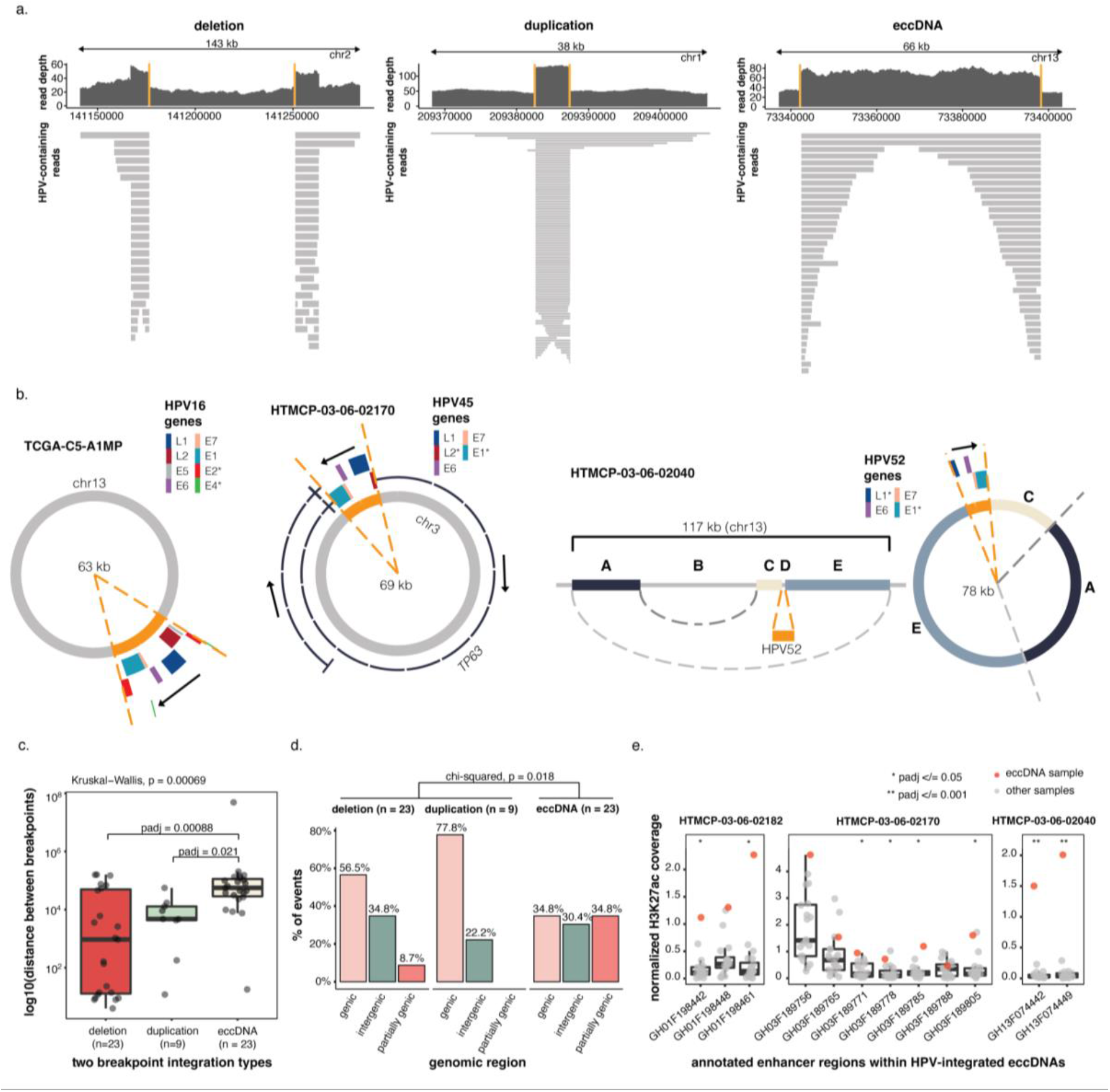
The three types of two-breakpoint integration events: deletion, duplication, and eccDNA. **(a)** Examples of read coverage patterns from deletion, duplication, and eccDNA integration events. Orange lines indicate the integration breakpoints. **(b)** Circular assemblies from three eccDNA integration events. The orange portions show the integrated HPV segment along with the integrated HPV genes. The direction of HPV gene transcription is shown by the black arrow. The right-most example shows a complex eccDNA event in which three non-adjacent human segments are shuffled together in the eccDNA. **(c)** The genomic distance between breakpoints in the three described integration event categories. **(d)** The percentage of events occurring in genic (>90% within a gene), intergenic (<10% within a gene), and partially genic (10-90% within a gene) regions, separated by integration category. **(e)** Normalized H3K27ac ChIP-seq coverage (RPKM) at GeneHancer-annotated enhancers contained on eccDNAs. Three HPV-integrated eccDNA-containing HTMCP samples with ChIP-seq data available are shown. The sample harboring the eccDNA integration is indicated by a red dot and compared to the rest of the cohort. Box plots represent the median and upper and lower quartiles of the distribution; whiskers represent the limits of the distribution (1.5 IQR below Q1 or 1.5 IQR above Q3). Adjusted *P* values in (c) were calculated post-hoc using a Benjamini-Hochberg-corrected Dunn’s test after a Kruskal−Wallis test. The *P* value in (d) was calculated using a chi-squared test. The *P* values in (e) were calculated using a permutation test and then were Benjamini-Hochberg corrected by sample. Where relevant, all statistical tests were two-sided.

The eccDNA integration events are unique because they exist outside the chromosome and form a circular structure bridged by HPV DNA (examples shown in Fig. 3b). The first example shows an HPV16 integration event that occurred in the 13q22 integration hotspot (Fig. 3b, left). Interestingly, 4/23 eccDNA events (two HPV16 integration events and two HPV45 integration events) occurred within this region, with all four breakpoint pairs overlapping each other, making this the only region recurrently involved in eccDNA events (Extended Data Fig. 3a). The second example shows an eccDNA event that contains a portion of *TP63* along with a portion of HPV45 (Fig. 3b, middle). The human and viral genes are predicted to be transcribed from opposite strands in this example, as HPV integrated in the inverse direction relative to *TP63*. The third example shows an atypical eccDNA integration event, in which the eccDNA contained a single HPV52 integrant along with a rearranged intergenic segment of chr13 (Fig. 3b, right).

The human-HPV breakpoints that comprised eccDNA integration events were distributed differently across the human genome compared to deletion and duplication events. Firstly, the spacing between the two breakpoints was greater and showed less variability in eccDNA integration events, with a median distance of 56 kb (Kruskal−Wallis, *p*=0.00069; Fig. 3c; Supplementary Table 6). The eccDNA integration events were also less likely to be completely contained within genic regions and more likely to have a breakpoint within the 3’ or 5’ end of a gene (chi-squared, *p*=0.018; Fig. 3d; Supplementary Table 6).

Previously generated H3K27ac chromatin immunoprecipitation followed by sequencing (ChIP-seq) data^19^ were available for five HTMCP samples with eccDNA events, allowing us to study the epigenetic status of annotated enhancers^20^ on these eccDNAs (Fig. 3e). The HPV45 (HTMCP-03-06-02170) and HPV52 (HTMCP-03-06-02040) integrated examples from Fig. 3b, as well as one HPV18-integrated sample (HTMCP-03-06-02182), contained enhancers that were significantly activated in the integrated sample compared to the rest of the cohort (Supplementary Table 7). Notably, the HPV52-integrated eccDNA event (HTMCP-03-06-02040) occurred in a region that appeared repressed in the cervical tumors in our cohort based on H3K27me3 enrichment, whereas the enhancers had H3K27ac coverage consistent with activation in the integrated sample (Fig. 3e). Interestingly, the HPV52 integrated sample featured the only integration event near *KLF12* associated with increased expression of *KLF12* relative to the other samples (Fig. 1g; top point in the box plot). In contrast to the other eccDNA events in this region (Extended Data Fig. 3a), the HPV52 eccDNA originated upstream (3’ direction) of the *KLF12* promoter. The two activated enhancers contained on the eccDNA (GH13F074442 and GH13F074449 from GeneHancer^20^) are known distal enhancers of *KLF12* (Fig. 3e). Human enhancers on HPV-integrated eccDNA may, therefore, function to activate both the eccDNA-contained HPV in *cis* as well as human genes on the chromosome in *trans*.

### HPV integration is associated with full chromosome arm translocations

Translocation integrations, in which two breakpoints occurred on different chromosomes, were rare in our study. There were only four instances of translocation integrations, involving four different HPV types (HPV16, 18, 52, and 31). Two examples of the read alignments from translocation integration events are shown in Extended Data Fig. 4a-b. The first example shows an HPV16-associated translocation event where integration breakpoints occurred in the pericentromeric regions of chromosome 4 and 12 (Extended Data Fig. 4a). The translocation contained a small (164 bp) segment of the HPV16 genome between the pericentromeric repeats on the chromosomes. The read depths indicated one copy loss in the p-arm of each chromosome immediately 5’ of the HPV breakpoints (Extended Data Fig. 4c), compatible with an unbalanced translocation in which the q-arms of chromosome 4 and 12 recombined via an HPV16 fragment and the p-arms were lost (Extended Data Fig. 4e). The second example, an HPV52-integrated translocation event, involved a focal amplification around the site of HPV integration and a segment of HPV that contained *E6* and *E7* (Extended Data Fig. 4b). The chromosome 8 breakpoint, although within a gene, occurred near the pericentromeric region and was adjacent to a one copy loss upstream of the breakpoint. The chromosome 1 breakpoint occurred in a genic region on the p-arm and showed copy gain across a ∼70 Mb segment upstream of the integration breakpoint (Extended Data Fig. 4d). This indicates that the chromosome 1 region was duplicated and recombined with the q-arm of chromosome 8, the p-arm of chromosome 8 was lost, and the HPV52 segment was focally amplified around the translocation junction (Extended Data Fig. 4f).

### Multi-breakpoint HPV integration events are structurally complex

Multi-breakpoint events, comprising 55/130 (42%) of all events, were the most common in our study. Fig. 4a summarizes the number of HPV-human breakpoints per multi-breakpoint event, which ranged from 3 to 32 (Supplementary Table 8). The majority of multi-breakpoint events (96%, 53/55) arose from HPV16/18 integrations and from one sample harboring multiple HPV58 integrations (Fig. 4a). On average, HPV18 integrations contained more breakpoints per event than HPV16 (Wilcoxon, 5.9 vs. 7.4, *p*=0.034; Fig. 4b), corresponding with our observation that HPV18 integrants had more viral genome copies than HPV16 integrants (Fig. 2i). Integrations with higher numbers of HPV breakpoints were also associated with more human-only structural variants (SVs) overlapping the integration event, indicating general instability within these regions (Spearman correlation, *R*=0.58, *p*=0.00037; Fig. 4c; Supplementary Table 8).

**Figure 4:**
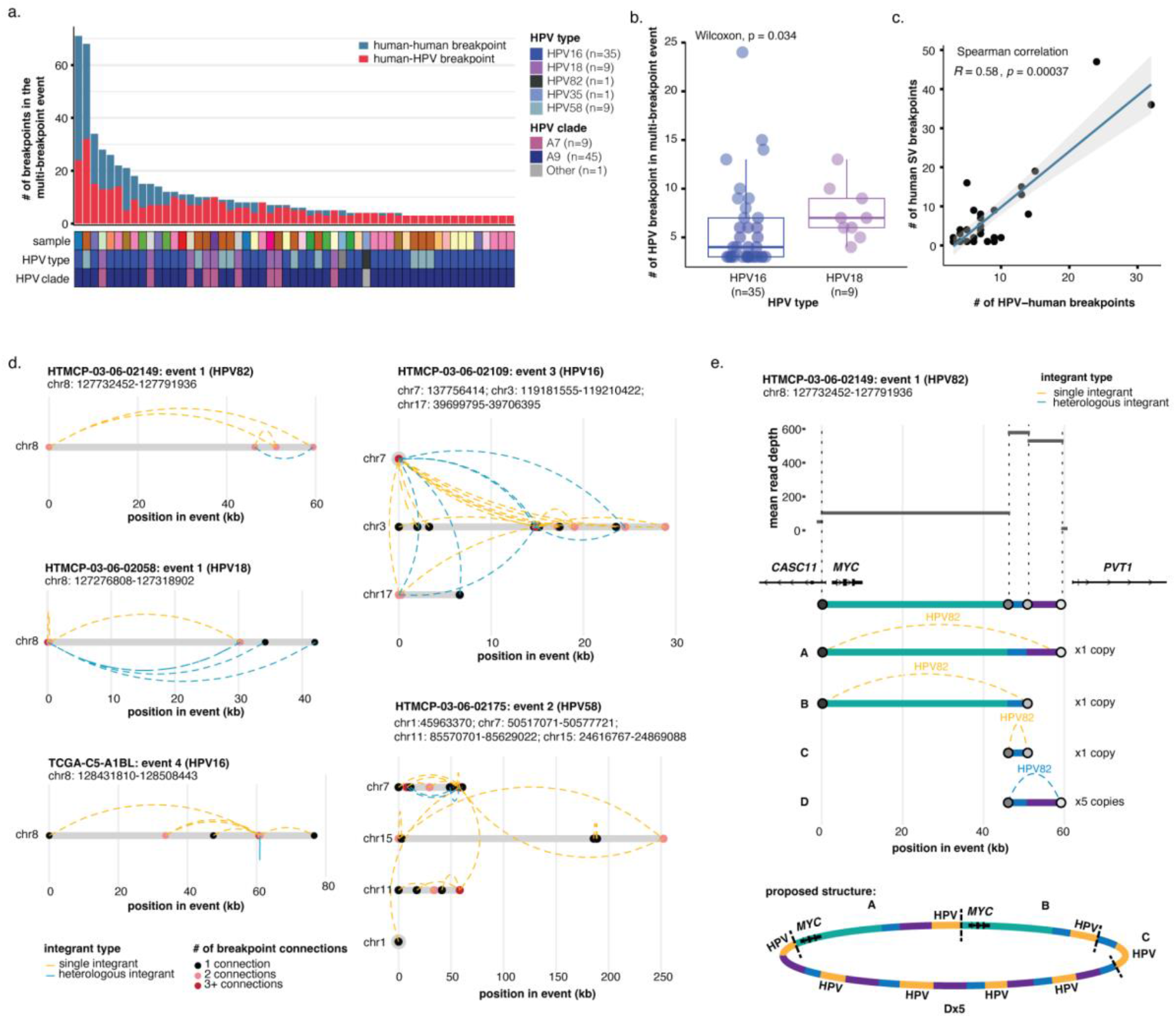
Complex structural variation overlaps multi-breakpoint integrations. **(a)** The number of human-HPV and human-human SV breakpoints across multi-breakpoint integration events, and the HPV type and clade in each. **(b)** The number of breakpoints per event in HPV16 and HPV18 multi-breakpoint events. **(c)** Spearman correlation between the number of HPV-human breakpoints and the number of human-human SV breakpoints in multi-breakpoint events. **(d)** Examples illustrating the connectivity between HPV breakpoint pairs in five multi-breakpoint events: three from the *MYC*-associated locus on chr8 and two from multi-chromosomal events. Dots denote HPV breakpoints along the loci, and dotted lines represent the HPV integrants that bridge the breakpoints. The breakpoints are colored according to how connected they are. The integrants are colored according to whether single (orange) or heterologous (blue) integrant structures bridge the breakpoints. **(e)** An example breaking down the copy number changes and the proposed structure of HTMCP-03-06-02149 event 1 overlapping *MYC*. Box plots represent the median and upper and lower quartiles of the distribution; whiskers represent the limits of the distribution (1.5 IQR below Q1 or 1.5 IQR above Q3). *P* values were calculated using the Wilcoxon rank-sum test and a Spearman correlation test.

We next determined how HPV breakpoints were connected within multi-breakpoint integration events. Five representative examples from the cohort are shown in Fig. 4d, three of which originate from the hotspot on chromosome 8 (Fig. 1f). The presence of heterologous integrants between breakpoint pairs is indicated, as well as the connectivity of each breakpoint (Fig. 4d). The three events from the chromosome 8 hotspot all span similar distances (40-80 kb) and contain similar numbers of breakpoints (4-6 breakpoints). HTMCP-03-06-02058 event one and TCGA-C5-A1BL event four both feature one centralized breakpoint that connects to all the other breakpoints in the event. The single centralized breakpoint was a recurrent theme across the multi-breakpoint events, especially in single-locus events. This contrasts with the HTMCP-03-06-02149 event, in which each breakpoint connects to two others, resulting in a continuous linked structure. The two multi-chromosome events shown in Fig. 4d were the two most rearranged events from Fig. 4a. The HTMCP-03-06-02109 event involved one heavily rearranged chromosome (chromosome 3), while chromosome 7 and 17 contained one and two breakpoints, respectively. Interestingly, the single breakpoint on chromosome 7 paired with almost every other breakpoint on the other two chromosomes. In contrast, HTMCP-03-06-02175 event two contained three highly-connected breakpoints (three or more connections) but none connected with all three other chromosomes. We hypothesize these patterns suggest two modes of generation: one catastrophic event that joins HPV concatemers to one central breakpoint (HTMCP-03-06-02109 event three), and joining together different HPV-integrated regions over the course of a tumor’s evolution (HTMCP-03-06-02175 event two).

To resolve the copy number patterns that can occur at multi-breakpoint events, we delved deeper into the structure of HTMCP-03-06-02149 event one (Fig. 4e). This event was of particular interest because it encompassed *MYC* and was predicted to be circular, based on the assembly and breakpoint connectivity. Read depth conversions were used to predict absolute copy number ratios for each segment bridged by a HPV breakpoint pair in the event. The three segments that were bridged by non-heterologous integrants, including the region containing *MYC*, each showed a one copy increase above the baseline adjacent to the event (Fig. 4e). The segment between the heterologous integrant breakpoint pair showed a five-copy increase, suggesting again that amplification may be involved in the generation of heterologous integrants (Fig. 4e). The resulting structure therefore contained 8 HPV integrants linking these human genome segments and two copies of *MYC* within a complex circular structure, as exemplified by the proposed structure in Fig. 4e.

### HPV integration events show consistent orientation-dependent methylation patterns

ONT sequencing simultaneously yields DNA methylation signals along with DNA sequence data^21^. We investigated methylation patterns around and within HPV integration, including methylation of the HPV genome. The methylation status of CpGs within HPV integrants and up to 5 kb on either side of the breakpoints is shown in Fig. 5a-b for eccDNA and deletion integration events (Supplementary Table 9 and Supplementary Table 10). The integration events are oriented by the direction of HPV transcription. Generally, we observed that human regions upstream of HPV transcription were methylated, while human regions downstream of HPV transcription were unmethylated. In all integrants, the long control region (LCR), a non-coding regulatory region that activates early HPV gene transcription^22^, was invariably hypomethylated compared to the genic region of HPV (Fig. 5a-e). In the eccDNA events, the late HPV genes (*L1* and *L2*) were methylated, while the early HPV genes (specifically *E6* and *E7*) were unmethylated following the LCR up to the 3’ breakpoint (Fig. 5a). The pattern of upstream hypermethylation and downstream hypomethylation on the human genome was consistent across most eccDNA integration events, although two HPV16 events showed uniform hypomethylation across the entire region (Fig. 5a; Supplementary Table 10).

**Figure 5:**
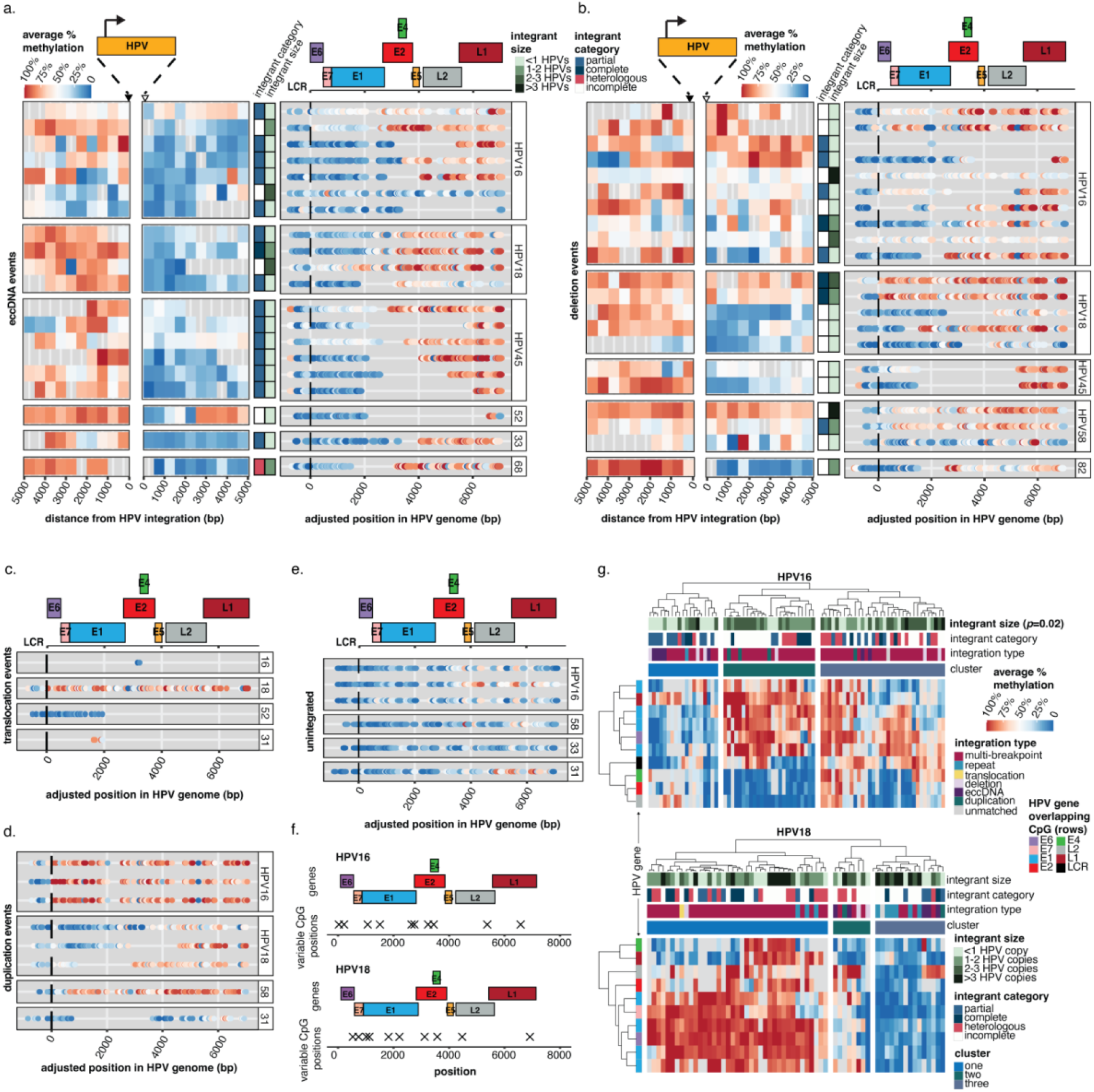
HPV integration causes distinct methylation patterns upstream, within, and downstream of the HPV genome. **(a,b)** The methylation frequency within and adjacent to HPV integrants in (a) eccDNA and (b) deletion integration events. The regions 5 kb upstream and downstream (relative to the direction of HPV transcription) are divided into 500 bp bins, and the average methylation frequency for all the CpGs within each bin is shown. Within HPV, the methylation of each CpG group is shown as a point. The integrant size and category are also indicated for each integrant (row). All the events are aligned to the start of the genic region (E6 start) for each respective HPV type. The gene model for HPV16 is shown above for general reference. **(c-e)** The HPV methylation frequencies for (c) translocation events, (d) duplication events, and (e) unintegrated HPV genomes as described in (a) and (b). **(f)** The positions of the top 10 most variable CpG positions in the integrated HPV16 and HPV18 genomes. **(g)** Hierarchical clustering of the HPV integrants using the 10 most variably methylated CpGs from (f). Clusters were determined using the ward D2 method with Euclidean distance measures. Integrant sizes in (g) were compared using the Kruskal−Wallis test, which was followed post hoc by a Benjamini-Hochberg corrected Dunn’s test; cluster one vs. cluster two, *padj*=0.024, cluster one vs. cluster three, *padj*=0.033.

Compared to eccDNA events, the other two-breakpoint integration types showed more variable methylation in the genic regions of HPV. Integration events that contained more than one copy of HPV often had inconsistent methylation between the read alignments, resulting in partial methylation or hypermethylation across the genic region (Fig. 5b-d). Deletion and duplication events containing partial copies of HPV showed similar patterning as the eccDNA events (Fig. 5a,b,d). Deletion events that were hypermethylated across the genic region of HPV also tended to show methylation in the flanking human regions (Fig. 5b). Three of the four translocation events contained small, partial HPV genome segments that were uniformly hyper- or hypomethylated across the segment, indicating that these events may not be regulated via the same mechanisms as the other integration types, and may instead take on the methylation pattern of the surrounding region (Fig. 5c).

The methylation patterns in the integrated HPV genomes contrasted with the six samples containing episomal HPV (Fig. 5e). Across all six episomal samples, the HPV sequences were less methylated than the integrated HPV genomes, particularly in the late genic region. A differential methylation analysis^23^ between the HPV16 integrants and the three episomal HPV16 genomes confirmed that methylation was significantly different across the majority of the HPV16 genome (positions 30-5125 and 5706-7858, DSS, *padj*<0.01; Supplementary Table 11). The non-significant region corresponded with the part of HPV (*E2*) that is often lost in the process of HPV integration^24^; therefore, less methylation data was available for this region in the integrated samples.

For HPV16 and HPV18 integrants, we subsequently determined the 10 most variable CpG sites to observe how methylation compares between integrants (Fig. 5f; Supplementary Table 12). These variable methylation sites were distributed equally across the HPV16 and HPV18 genomes, with a small cluster of variable CpGs in the *E1*/*E2* genes of HPV16 (Fig. 5f). Notably, one of the HPV16-associated variable CpGs was located at position 30-57 in the LCR, which overlaps with the SP1 and E2 binding sites 3 and 4 (E2BS3 and E2BS4)^25^. We used the variable methylation sites to cluster the HPV16 and HPV18 integrants and identified three methylation clusters each for HPV16 and HPV18 integrants (Fig. 5g; Supplementary Table 13 and Supplementary Table 14). In the HPV16 CpGs, cluster two integrants were consistently methylated at variable CpGs located in *E1*/*E2*/*E6*, while cluster one was generally less methylated and cluster three was partially methylated across all positions (Fig. 5g; top). The HPV16-integrated eccDNA and deletion events were mostly grouped in cluster one and cluster three, respectively (Fig. 5g; top). The CpGs in HPV18 separated integrants into three clusters that were methylated (cluster one), partially methylated (cluster two), and unmethylated (cluster three) (Fig. 5g; bottom). Interestingly, most repeat region integrants grouped with the unmethylated cluster (cluster three), and eccDNA integrants were split between clusters two and three (Fig. 5g; bottom). Cluster one was predominantly composed of multi-breakpoint integrants, indicating that these events harbored more DNA methylation at the variable CpGs (Fig. 5g; bottom). No significant clustering of the integrant categories was observed, although this may be due to the number of incomplete integrants represented. HPV16 integrants in cluster one (unmethylated cluster) were significantly shorter than integrants in clusters two and three (Kruskal−Wallis, *p*=0.02; Benjamini-Hochberg corrected Dunn’s test, cluster one vs. cluster two, *padj*=0.024, cluster one vs. cluster three, *padj*=0.033). Integrant sizes were not significantly different between HPV18 methylation clusters.

### Allele-specific expression and methylation patterns in regions of HPV integration

HPV integration affects only one allele. We leveraged this to explore how HPV integration may have allele-specific impacts on the epigenome and human gene targets on a broader scale. We phased reads into haplotypes and determined the allele from which the HPV-containing reads originated, and then identified differentially methylated regions (DMRs) between the two haplotypes across the tumor genome (Methods). DMR density was compared to a null distribution across the human chromosomes to identify regions with significantly dense clusters of DMRs, referred to as DMR hotspots (Methods). HPV integration sites were then intersected with the DMR hotspots to identify events that potentially affected DNA methylation in an allele-specific manner, which would suggest HPV-associated epigenomic dysregulation. Across the samples in our dataset, 27 integration event loci overlapped autosomal DMR hotspots (Fig. 6a; Extended Data Fig. 5a-e; Supplementary Table 15). HPV-overlapping DMR hotspots had a median size of 5.4 Mb (ranging from 1.9 Mb to 26 Mb) compared to the overall mean of 3.6 Mb (ranging from 18 kb to 38 Mb) and mostly encompassed multi-breakpoint events (Extended Data Fig. 6a-b).

**Figure 6:**
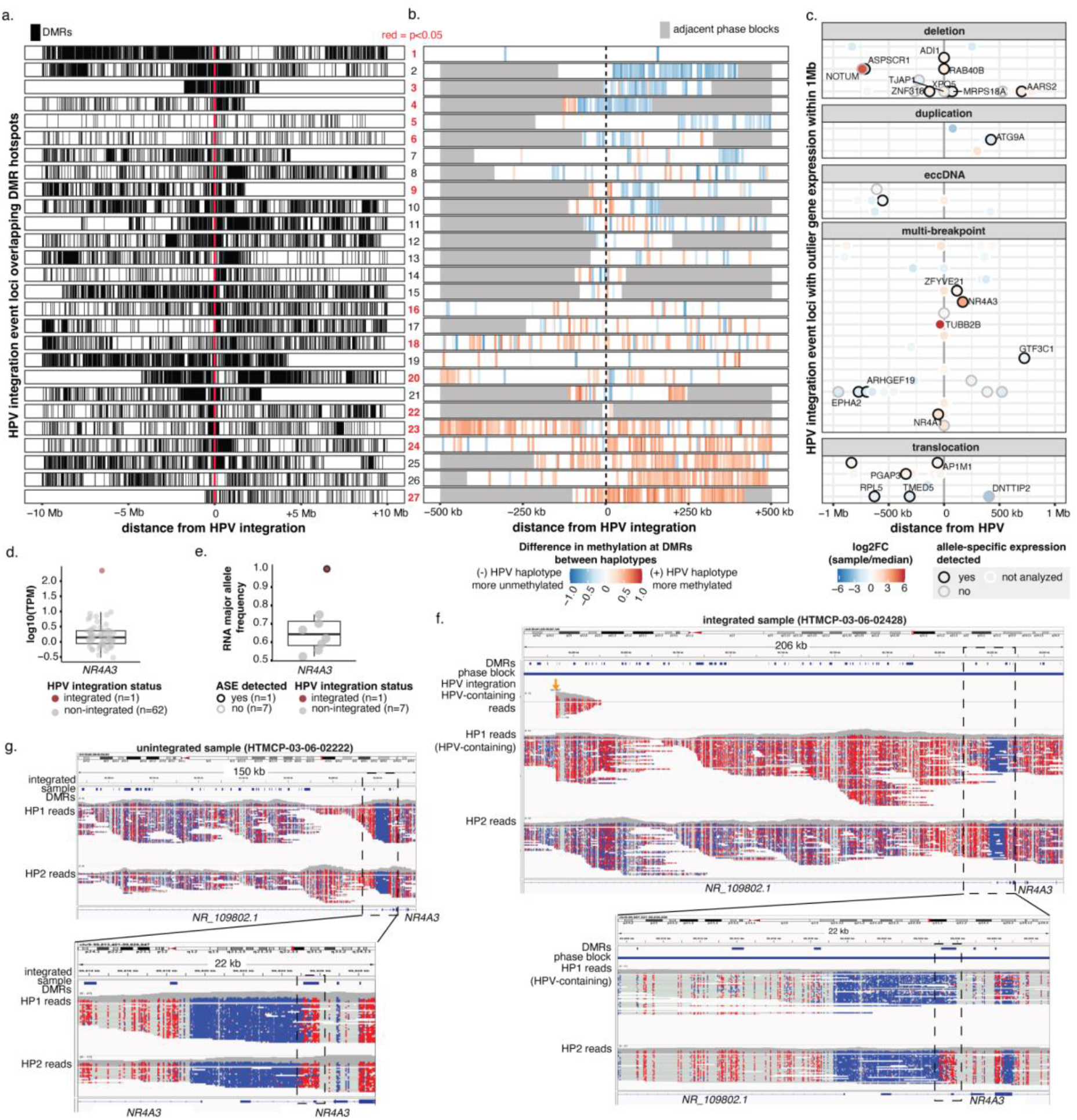
HPV dysregulates the methylome on the integrated haplotype. **(a)** The distribution of DMRs across 10 Mb on either side of HPV integrations in the 27 events that overlap a DMR hotspot. **(b)** The direction of methylation changes in the HPV-containing haplotype within the phase block containing HPV integration. Adjacent phase blocks are shown as flanking gray bars. Events are ordered identically in (a) and (b). The event numbers are colored red when *p*<0.05, as determined by a permutation test. **(c)** The position of genes with outlier expression (1.5 IQR below Q1 or 1.5 IQR above Q3) relative to sites of HPV integration. Expression fold change in the integrated sample relative to the median of the cohort and ASE status are also indicated. **(d)** The expression of *NR4A3* in the integrated sample (HTMCP-03-06-02428) compared to other samples in the dataset. **(e)** The RNA-seq major allele frequency and ASE of *NR4A3* in HTMCP-03-06-02428 compared to other samples in the dataset. **(f)** Integrative Genome Browser snapshots showing wide (top) and zoomed (bottom) views of the haplotype-specific methylation changes around *NR4A3* and HPV integration in HTMCP-03-06-02428, with reads separated into the two haplotypes (HP1 and HP2). The sample’s DMRs, phase blocks, and HPV integration breakpoints are also indicated in the top three tracks. **(g)** Integrative Genome Browser snapshots showing wide (top) and zoomed (bottom) views of methylation around *NR4A3* in an unintegrated sample (HTMCP-03-06-02222). HTMCP-03-06-02428’s DMRs are indicated in the top track. Reads in both (g) and (h) are colored by CpG methylation status, with red indicating methylated and blue indicating unmethylated.

We further investigated methylation around the 27 HPV integration events overlapping DMR hotspots (Fig. 6a-b). Among DMRs within the phase block that contained the HPV integration, we observed variation in DMR distribution, in regards to their position relative to HPV and in size and density. For example, event three was centered on a dense, 5 Mb hotspot with few DMRs outside the hotspot (Fig. 6a). In contrast, events 14, 17, 22, and 24 were flanked on one side by 500 kb – 1 Mb of dense DMRs and a reduced density of DMRs further away from the integration event (Fig. 6a). We also investigated the direction of DNA methylation changes on the integrated haplotype (Fig. 6b). In seven events (events 1-7), the DMRs were predominantly hypomethylated on the HPV-integrated allele, and eight events (events 20-27) showed hypermethylation on the HPV-containing allele, while the remaining events (events 8-19) did not have a discernible pattern (Fig. 6b; Supplementary Table 15, examples of large-scale HPV-associated methylation around integration events are shown in Extended Data Fig. 7 and Extended Data Fig. 8). Lastly, we tested if the DMR hotspots were a unique occurrence at that location in the integrated sample and not a result of the integration taking place in a commonly-occurring DMR hotspot across all samples (Methods). Eleven of the integration events showed significantly high (*p*<0.05, permutation test) DMR density at the overlapping hotspot compared to 1000 sampled regions, suggesting that these DMRs may be caused by HPV integration, while the others may represent instances of HPV becoming integrated in pre-existing DMR hotspots of the genome (Fig. 6a-b; Extended Data Fig. 6c).

We searched for nearby genes that may be transcriptionally affected by HPV integration, limiting our analysis to genes within 1 Mb of HPV integration sites. 83 genes were outliers, showing expression levels that were more than 1.5 times the interquartile range (IQR) above the third quartile (Q3) or below the first quartile (Q1) in 16 integrated samples and 32 integration event loci (Fig. 6c; Supplementary Table 16; Extended Data Fig. 6d). We examined these genes for allele-specific expression (ASE), which would suggest a *cis*-acting element (i.e. HPV integration) was altering gene expression on the affected allele. Of the outlier genes that we could test for allelic ratios (Methods), we found 24 (65%) that showed ASE (Supplementary Table 16). Nuclear receptors *NR4A1* and *NR4A3*^26^ were the genes most highly overexpressed in an allele-specific manner from the HPV-containing haplotype (Extended Data Fig. 6d-e). *NR4A1* (chromosome 12) and *NR4A3* (chromosome 9) were detected from two independent integration events in two different samples. Outlier expression and ASE of *NR4A3* was only detected in the integrated sample and was associated with a DMR hotspot (Fig. 6d-e), while *NR4A1* showed allele-specific expression in nine other samples (Extended Data Fig. 6e).

To determine how HPV may contribute to the upregulation of oncogenes, we sought to characterize the DNA methylation landscape around the HPV integration in the *NR4A3*-activated sample (HTMCP-03-06-02428). The integration event was a multi-breakpoint event, with one breakpoint on chromosome 9 upstream of *NR4A3* that bridged to a chromosome 14 locus harboring twelve breakpoints (Extended Data Fig. 6f). The chromosome 9 locus overlapped a DMR hotspot that was hypermethylated on the integrated allele (Fig. 6f, top). However, several DMRs near the *NR4A3* promoter were unmethylated on the integrated allele. Most notable was an 821-bp DMR that overlapped exon two of *NR4A3* at the 3’ end of an unmethylated promoter element (Fig. 6f, bottom). The HPV-containing allele was demethylated in this DMR, a feature that was absent in other cervical cancers we profiled (Fig. 6g; bottom). The broad methylation seen on the HPV-containing allele was also absent in other samples (Fig. 6g; top). We thus observed, in the *NR4A3*-activated sample, HPV-associated large-scale allelic methylation with targeted demethylation of *NR4A3* regulatory elements.

## Discussion

HPV integration plays a crucial role in the development of HPV-driven cancers. However, its effect on the structure and regulation of the genome remains incompletely understood due in large part to the complex genomic dysregulation associated with HPV integration. We used ONT long-read sequencing to achieve read lengths that could span from one HPV-human breakpoint to another, thereby enabling reconstruction of complex events. Our study thus aimed to provide a detailed overview of genomic structures and epigenomic changes associated with HPV integration events in cervical cancer and encompassing various HPV types. Based on the alignment patterns of HPV-containing reads, we then created a workflow to categorize and describe HPV integration events according to their distinct features on the HPV and human genomes. This study presents the most detailed characterization of HPV integration structures to date and is the first to relate specific HPV-associated structural alterations to local (+/− 5 kb) and distal (+/− 10 Mb) epigenomic changes in the tumor.

The two-breakpoint HPV integration events included deletion, eccDNA, duplication, and, less often, translocation events. Repeat region integration events occurred within short tandem repeat elements such as telomeres and centromeres, but the exact number of breakpoints and structures could not be accurately determined due to limited resolution in highly repetitive regions. The eccDNA events showed the most consistent methylation patterns on the HPV DNA and adjacent human DNA, with methylation in the human region upstream of the first breakpoint persisting on the HPV DNA, and the hypomethylated state beginning at the non-coding LCR region continuing through the second breakpoint to downstream human DNA. This pattern was also observed in a subset of the deletion integration events, but much more heterogeneity was present in events residing on the human genome. HPV eccDNAs can exist as self-contained regulatory systems that function to upregulate their embedded genes^27^, while intrachromosomal events are subjected to positional effects of the neighboring host chromatin. There was a genomic hotspot of eccDNA integration in the intergenic region between *KLF5* and *KLF12* on chromosome 13, but in contrast with the *MYC* hotspot, expression of *KLF5*/*KLF12* was not significantly changed in the integrated samples. From this we infer that these integration hotspots may arise due to selection for different attributes, for example, for events that drive *MYC* expression or events that excise regulatory elements between *KLF5* and *KLF12*. The four eccDNAs from this hotspot harbored nearly-overlapping non-coding regions of human DNA, suggesting that integration within non-coding elements may serve to drive HPV gene expression rather than amplify a human oncogene. In eccDNA samples for which ChIP-seq data were available, we found activation of human oncogene enhancers on the eccDNA, thus supporting the notion that HPV integration may select for attributes that provide a proliferative advantage.

In 24% of HPV integrants, the supporting reads indicated the existence of different integrant sizes between the same two breakpoints, which usually differed from each other by multiples of the HPV genome. The phenomenon of heterologous integration was first suggested by Kadaja et al.^15^, who showed that, upon integration, E1 and E2 drive re-replication of the HPV genome, generating intermediate structures whose resolution by DNA damage repair machinery leaves several heterologous structures. In our samples, heterologous integrants were only found in regions with amplification between the breakpoints (i.e. they were not found in deletion events). This supports Kadaja et al.’s observations and extends them by suggesting that the heterologous integrants represent different copies within the amplification rather than different cell populations in a heterogeneous tumor. We also found integrants with several tandemly amplified HPV genomes. Integrants containing multiple copies of HPV showed stronger methylation of the HPV genic regions. We thus deduce that tandem copies within a single amplified integrant could exist in different methylation states, thus explaining discordant methylation signals between supplementary alignments from the same read.

Of the six integration types identified, multi-breakpoint integration events were the most prevalent. These events contained between 3 to 33 HPV-human breakpoints across one or more loci and were associated with other SVs in the tumor genome. We reconstructed the HPV integration events from the two most complex multi-breakpoint events in our cohort, which involved three and four different chromosomes, respectively. One integration event had a single breakpoint "hub" that connected to most other breakpoints in the event, while the other involved three highly-connected breakpoints that joined different regions of the event. These patterns may suggest different mechanisms. The single centralized breakpoint suggests that the event arose at a single time point in a highly fragmented region, which was then reassembled with multiple HPV integrants linking the fragments, akin to a local chromothripsis event^28^. Akagi et al.^29^ described this patterning of HPV integration events as “heterocateny”, or a “variable-chain”, where several concatemers of bridged human and HPV segments are duplicated but with new breakpoints arising between duplicated copies. In contrast, the event with multiple highly-connected segments suggests that the events may have gained complexity throughout the tumor’s evolution. When new double-stranded breaks occur, HPV integrants scattered across the genome may enable recombination between these regions through shared homology.

HPV integrants within multi-breakpoint events were commonly heterologous and methylated, and associated with altered expression of nearby genes. The broad methylation of the HPV-integrated allele could arise as a host defense mechanism in response to viral integration, as reported for other viruses and endogenous retroviruses^30–32^. We identified a genomic hotspot of multi-breakpoint integration events near the *MYC* locus on chromosome 8, which was associated with increased *MYC* expression in the integrated samples. Chromosome loops in this region that connect the integrated locus to *MYC* in a haplotype-specific manner have been described in HeLa cells^10^. We also observed HPV-associated activation of *NR4A3*, which was situated 170 kb downstream (3’) of an HPV breakpoint that was part of a multi-breakpoint event bridging two chromosomes. *NR4A3* is a nuclear receptor and transcription factor with both oncogenic and tumor suppressive roles documented in cancer^33–36^. It is also a transcriptional target of p53, which is known to be targeted for degradation by E6 in HPV-driven tumors, suggesting its expression would normally be reduced^37,38^. A rare tumor, salivary gland acinic cell carcinoma, is driven by translocation events that juxtapose super-enhancers from other chromosomes to the upstream regions of *NR4A3*^39,40^. We speculate that the HPV-driven activation of *NR4A3* may occur through a similar mechanism. The ASE of *NR4A3* in this sample and its distance from the HPV breakpoint also suggests a physical interaction between enhancer elements involved in the HPV integration event and *NR4A3.* These enhancers may come from the HPV LCR or from the bridged chromosome. Further studies are required to discern how HPV integration leads to activation of certain genes while driving megabase-scale hypermethylation on the affected haplotype.

Overall, our study identified patterns in the structure and regulation of human and viral elements of HPV integration events. We observed methylation and expression changes in *cis* with the integration event. Further studies could reveal underlying mechanisms that cause HPV integration, how and if those mechanisms result in the different genomic structures we observed, and the extent to which positional effects of human genome neighborhoods (e.g. euchromatin and heterochromatin) and resultant methylation changes influence epigenetic dysregulation around HPV integration events.

## Acknowledgements

We are grateful for contributions from the other members of various groups at Canada’s Michael Smith Genome Sciences Centre, including those from the Biospecimen, Library Construction, Sequencing, Bioinformatics, Technology Development, Quality Assurance, LIMS, Purchasing and Project Management teams. V.L.P. and L.C. were the recipients of CIHR Frederick Banting and Charles Best Canada Graduate Scholarships GSD-152374, GSD-164207 respectively. M.N. was supported by postdoctoral fellowship awards from the Canadian Institutes of Health Research (FRN-188098) and Micheal Smith Health Research BC (RT-2023-3168). S.J.M.J. is the recipient of the Canada Research Chair in Computational Genomics. M.A.M. is the recipient of the Canada Research Chair in Genome Science. This work was supported in part by funding provided by the Canadian Institutes for Health Research (CIHR award FDN-143288 and PJT-180410) to M.A.M. and NCI R21CA241013 and R01CA262198 to J.S.R. and MCW OBGYN WHRP fund.

## Author Contributions

This project was conceived by V.L.P. and M.A.M. Data were generated by Canada’s Michael Smith Genome Sciences Centre at BC Cancer and analyses were performed by V.L.P., K.O., and S.M. M.A.M. supervised this work. Contribution to methods used in data analyses: V.L.P., K.O., R.C., L.C., Z.H., G.C., J.F., K.M.N., V.A., I.B., and S.J.M.J. Contribution to data generation: K.O., R.C., M.I., R.M., S-W.T., S.K.C., J.H., R.M., E.C., K.L.M., A.J.M., S.J.M.J., and J.S.R. V.L.P., M.A.M, and M.N. wrote the manuscript. All authors reviewed and edited the manuscript.

## Code Availability

All code used for analyzing HPV integration events using ONT long-read WGS and Illumina short-read WGS can be found as an open source workflows on GitHub (ONT: https://github.com/vanessa-porter/callONTIntegration, Illumina: https://github.com/vanessa-porter/illuminaCallHPVInt). The code used to make the figures in this paper can also be found on GitHub (https://github.com/vanessa-porter/Porter-ONT-HPV-FigureScripts2023).

## Data Availability

The raw long-read sequencing data, DNA methylation calls, SV calls, SNV calls, and phasing results will be deposited to an accessible and controlled database. Other data from the HTMCP dataset used in this study have been deposited in dbGaP under the dbGaP study ID (<u>phs000528</u>) as part of the NCI Cancer Genome Characterization Initiative (CGCI; <u>phs000235</u>). The processed expression data used in this study can be downloaded from the Genomic Data Commons (GDC) data portal (https://portal.gdc.cancer.gov/).

## Methods

### Sample selection

Our sample selection balanced HPV types, homologous recombination deficiency (HRD) scores (which may impact SV generation)^41^, and HPV integration statuses, to what was previously identified across the whole cohort using short-read sequencing^18,19^. Our final dataset included 43 samples from the HTMCP^19^ and 20 samples from TCGA^18^, which included 26 HPV16 tumors, 17 HPV18 tumors, 8 HPV45 tumors, and 12 tumors with less common HPV types, approximately reflecting the proportions of HPV types present in the larger TCGA and HTMCP cohorts. Visual inspection of the HPV genome alignments was performed to confirm the HPV-containing reads were of high-quality, both in mapping quality and in length and contiguity. Eleven samples (one HTMCP, ten TCGA) that were initially sequenced were excluded in the final dataset due to an insufficient number of HPV-aligning reads, ambiguous or low-quality alignments, or highly fragmented and uninterpretable reads across the HPV genome due to low n50s and high chimeric rates in the sample. None of these samples had HPV integration detected by our workflow, but their low-quality alignments disqualified them from accurately representing episomal HPV.

### ONT library construction and whole-genome sequencing

For TCGA samples, DNA was re-extracted from fresh frozen tissue. 14 HTMCP samples had sufficient DNA left over from an earlier extraction^19^ but 29 samples required fresh extraction. For archival (frozen) tissues, nucleic acids were extracted using bead-based or column purification methods. Blue Pippin size-selection (Sage Science) was performed on 5 µg DNA to deplete shorter DNA molecules (<15 kb) from the final library to achieve 2 µg of final input DNA at a concentration of 42 ng/µL. The ONT ligation-based library preparation kit (SQK-LSK110) was implemented on a NIMBUS liquid handling robot (Hamilton), followed by the whole genome PCR-free library construction for nanopore sequencing. The libraries were loaded onto quality-controlled flow cells exhibiting >5000 active pores within 5 days of construction to preserve the pore-targeting adapter. The library recovery after ligation and bead purifications was expected to be >40% of input (i.e. >800 ng). For yields 300-500 ng the entire library was loaded onto a PromethION flow cell. For libraries with >500 ng yield, 2/3 of the library was loaded initially followed by a nuclease (DNase I) flush of the flow cell to restore the activity of the pores that had become clogged with DNA. The remaining 1/3 of the library was then loaded onto the restored flow cell for a maximum total of 750 ng. PromethION flow cells were typically run for 72 hours for maximal sequencing yield.

### ONT Primary Data Analysis

Raw signal from the PromethION sequencer was basecalled using ONT’s Guppy 5, with the “super-accurate” model. These sequence data were aligned using minimap (v. 2.15) to a custom reference containing GRCh38 with no ALT contigs and 17 HPV strains from the HTMCP and TCGA cohorts. Subsequent primary analysis was carried out via a NextFlow workflow. Structural variants were called from the aligned BAM using Sniffles^42^ (v. 1.0.12b) and CuteSV^43^ (v. 1.0.12). Small variants were called using Clair3^44^ (v. 0.1-r8) and phased using WhatsHap^45^ (v. 1.0), with phase blocks retained for later analyses. DNA methylation (5-mC) was called at the read level using Nanopolish^21^ (v. 0.13.2). The DNA methylation calls were phased into haplotypes using NanoMethPhase^17^ (v. 1.0). Methylation calling failed on one TCGA sample (TCGA-C5-A2LX) and one HTMCP sample (HTMCP-03-06-02176) and methylation phasing failed on six TCGA samples (TCGA-C5-A2LX, TCGA-C5-A1MN, TCGA-C5-A2LY, TCGA-C5-A3HE, TCGA-C5-A7X3, and TCGA-C5-A8YR) and one HTMCP sample (HTMCP-03-06-02176).

### Identifying HPV integration breakpoints, integration events, and integration event loci

Sniffles (v. 1.0.12) was used to call translocations on the ONT long-read WGS using the following specifications to maximize the accuracy for detected HPV integration breakpoints: max_distance = 50, max_num_splits = −1, report_BND, num_reads_report = −1, min_support = 5, and min_seq_size = 500. A minimum of 5 consensus reads was required to detect an HPV integration breakpoint. An R script (v. 4.0) was developed that then iteratively grouped together HPV breakpoints if they had one or more reads that overlapped between the breakpoints as indicated in the VCF. A distance threshold of 500 kb was also implemented using BEDTools (v. 2.30.0) to further group breakpoints that mapped near each other but may have lacked reads long enough to intersect. The first condition ensured that breakpoints appearing distant to each other relative to the reference genome but were physically linked through fusion rearrangements, could be paired together. The second condition ensured breakpoints mapping near each other but lacked a read long enough to link them together. The collection of HPV breakpoints that were grouped together through these two methods were referred to as an integration event. All read names belonging to an integration event were retained for later analyses. Integration event loci were defined as the integration breakpoints within an event that map within 500 kb of each other, as determined using BEDTools (v. 2.30.0). Therefore, integration events spanning multiple chromosomes or large genomic expanses would have multiple integration event loci. The integration event loci were used for regional analyses such as determining neighboring genes.

### HPV integration comparison between ONT and Illumina

The WGS data for the HTMCP fresh frozen samples were realigned to the hg38/HPVs reference genome using minimap2 (v. 2.15). The short-read SV caller Manta^46^ (v. 1.6.0) was used to call translocations between the human chromosomes and HPV genomes, and a minimum congruent read pair threshold of five reads was filtered with bcftools (v. 1.15.1). Integration breakpoints within 500 kb of each other were combined into integration events using BEDTools (v. 2.30.0). Subsequently, integration breakpoint calls for 41 HTMCP samples (samples with integration calls in one or both technologies) were summed and compared by sample and by event using tidyverse^47^ (v. 1.3.1, R) and ggpubr^48^ (v. 0.4.0, R) functions. The genomic region overlap of integration breakpoint calls for Illumina and ONT was determined using the BEDTools (v. 2.27.1) intersect function against hg38 annotations obtained from the Table Browser page on the University of California, Santa Cruz website^49^. Specifically, the GENCODE V43 knownGene annotation filtered to exons^50^, cpgIslandExt, and RepeatMasker^51^ annotations were downloaded as bed files (accessed May 2, 2023). Remaining integration breakpoint calls were classified as intronic/intergenic.

### Assembling HPV integration event contigs

Each HPV integration event was assembled into an integration contig for characterization. The events were first subsetted into event-only bam files with Picard tools’ (v. 2.26.6) FilterSamReads function using the read name text files created when grouping the integration events. The reads were then converted into FASTQ files using SAMtools (v. 1.12). Each set of event reads were then run through the assembler Flye^52^ (v. 2.9) with three rounds of polishing. The assembly was mapped back to the reference chromosome for assembly annotation and the reads were mapped back to the assembly to check assembly quality using minimap2^53^ (v. 2.23). Sniffles^42^ (v. 1.0.12) was run on the reads aligned to the assembly to check for rearrangements that may not be assembled correctly such as insertions, deletions, and duplications.

### Circular multi-breakpoint and atypical eccDNA detection

Event circularity was tested on all events to discover unconventional eccDNA events that did not follow the typical pattern but still were predicted to be extrachromosomal. The assemblies of events that successfully assembled were aligned to the reference genome using minimap2^53^ (v. 2.23). The alignments of the event reads were then subtracted using BEDTools (v. 2.30.0) from the assembly alignment. If there was minimal to no coverage outside the assembly region and there was adequate coverage (>2 reads) on each border of the assembly, then circularity was predicted. Extra confidence was given to assemblies that contained a single contig and were predicted as circular by Flye^52^ (v. 2.9).

### Categorizing HPV integration events

The categorization of the integration events followed the decision chart in Fig. 1c. Repeats were detected on the fasta files of the reads using RepeatMasker^51^ (v. 4.2.1). The output GFF file was then checked for the following repeat sequences: TAR1, ALR, HSAT4, HSAT5, HSAT6, and SST1. If over 50% of the reads contained repeat sequences, then the integration event was categorized as a repeat integration. If no repeat was detected, then the number of breakpoints in the event was counted. Events with more than two breakpoints were categorized as multi-breakpoint events and noted as circular if circularity was predicted. Two-breakpoint events were then categorized as translocation integration if the two breakpoints were on different chromosomes and deletion integration if the region in between the two breakpoints had no read coverage in the HPV-containing reads. The eccDNA integration and duplication integration were then differentiated by the HPV-containing read coverage outside the two breakpoints; duplication integration had read coverage outside the breakpoints, while eccDNA integration had none. These conditions were measured and tested using a series of python scripts that categorized read coverage as enumerated by BEDTools (v. 2.30.0).

### HPV integrant copy number calling

The HPV-containing read alignments in PAF format were analyzed using an R (v. 4.0.2) script to determine the HPV integrant lengths in a read-specific manner. Reads containing the same human and HPV breakpoints were grouped together, allowing a leniency in the exact breakpoints of +/− 30 bp. The HPV alignments that occurred between two SV-detected breakpoints were extracted from each read and the total length of the HPV integrant was determined by its cumulative length on the read. If only one breakpoint was detected, the length of the HPV alignments before or after the break was calculated for incomplete HPV integrant analyses. All unique breakpoint pairs (or incomplete breakpoints) were determined for each sample. The HPV integrant sizes for each breakpoint pair were separated into read groups if there was a difference of >300 bp to the next closest size. Reads with integrant sizes within <300 bp of each other were therefore grouped together. The size of each unique breakpoint pair or breakpoint was the categorized: “heterologous” if >1 integrant size group was detected, “partial” if the HPV integrant was <1 HPV genome equivalent in size, “full” if the HPV integrant was >1 HPV genome equivalent in size, and “incomplete” if the breakpoint was not paired. The maximum integrant size of heterologous integrants was determined as the mean size of the largest read group. The maximum size of incomplete integrants was determined by the read containing the longest HPV alignment before or after the breakpoint.

### Two-breakpoint event comparisons

The genomic distance between breakpoints was calculated for all two-breakpoint events by subtracting the reference position of the most 5’ breakpoint from the most 3’ breakpoint. A Kruskal−Wallis test was performed on the three groups and was followed post-hoc by a Benjamini-Hochberg corrected Dunn’s test. The position of the two-breakpoint events relative to genic regions was determined by intersecting the region between the two-breakpoint (event region) with the gene regions (Ensembl 100 GRCh38) using BEDTools (v. 2.30.0) ^54^. The events were categorized as “genic” if >90% of the event region fell within a gene, “partially genic” if the event was between 10-90% within a gene, and “intergenic” if <10% of the event region was within a gene. The proportions of events that were genic, partially genic, and intergenic were compared between the three two-breakpoint event categories using a chi-squared test.

### GeneHancer H3K27ac coverage in eccDNA events

Five HTMCP samples with eccDNA events also had ChIP-sequencing available^19^. We leveraged this to compare the enhancer activation of the sample with the eccDNA to samples that do not have eccDNA in that region. The normalized coverage of H3K427ac (reads per kilobase per million, RPKM) was calculated in all HTMCP samples within our dataset with ChIP-seq available (n=20) at GeneHancer annotated enhancer regions that intersected with eccDNA regions using deepTools (v. 3.0.2)^55^ and BEDTools (v. 2.30.0)^54^, respectively. The fold change difference between the normalized coverage of the eccDNA-containing sample and the mean normalized coverage of all other samples was compared to a random sampling of 1000 enhancer regions. The *P* value was determined by the number of randomly selected enhancers that had a greater fold change in the eccDNA sample than the tested enhancer region. *P* values were then adjusted using a Benjamini-Hochberg correction.

### Copy number analysis

The copy number profiles were obtained for TCGA-A5-A1BL and HTMCP-03-06-02054 and other translocation event-containing samples. The Ploidetect^56^ (v. 3.0) pipeline was ran using the “short” sequence type for HTMCP samples and the “ont” sequence type for TCGA samples since high-coverage short-read WGS was unavailable (see https://github.com/lculibrk/Ploidetect-pipeline, section 1.1.4). The results were selected for the first predicted model that did not have a ploidy of 1.

### Multi-breakpoint event reconstruction and visualization

The multi-breakpoint event loci were visualized using a custom R script that positioned the HPV integrant connections. The HPV breakpoint pairs were grouped into their respective events, and the selected multi-breakpoint events in Fig. 4d were visualized by connecting the two breakpoints within a breakpoint pair using a dotted arched line. The heterologous integrants were differently colored to the integrants with a single event structure. The number of connections a single breakpoint had (i.e. the number of breakpoint pairs it was in) was also indicated by the color of the dot. Read depth of HTMCP-03-06-02054 event 1 was calculated using SAMtools (v. 1.12)^57^ in the regions between each breakpoint.

### HPV genome methylation and flanking methylation

The HPV genome methylation frequencies per CpG group were calculated using the methylation outputs from Nanopolish^21^ (v. 0.13.2) on the HPV-containing reads from each group of reads belonging to a single HPV integrant (i.e. a unique breakpoint pair). Those reads were also used to calculate the average methylation flanking the integrant for the deletion and eccDNA integration events. For the flanking regions, the methylation frequencies were averaged across 500 bp bins up to 5000 bp upstream and downstream of the breakpoints on the human DNA. The upstream/downstream directions were oriented with the strandedness of the HPV integrant so that the directions were relative to the direction of transcription on the HPV genome. For visualization, the HPV genome positions were shifted to begin at the *E6* start position and end at the *L1* stop position, with all LCR positions represented as negative values so they could be visualized together.

### Variable HPV methylation clustering analysis

The variation of the methylation frequencies for each CpG probes and/or clusters were calculated across all HPV integrants for HPV16 and HPV16 (excluding NAs). The positions were then ranked by variability and the top 10 positions were selected for both HPV types. The HPV integrants were then clustered and visualized using pheatmap^58^ using the ‘ward.D2’ method with ‘euclidean’ distances. Integrants that did not exhibit enough variability in the CpG positions to calculate distance matrices were excluded from the clustering.

### Hotspots of differentially methylated regions

Differentially methylated regions (DMRs) were identified using the DSS^23^ 2-sample analysis. DMR hotspots were identified using an R script that compared the actual density of DMRs across each chromosome to a Guassian null distribution of the same number of DMRs. DMR hotspots were defined as regions where the actual density of DMRs was significantly higher than the null distribution. These hotspots were intersected with the HPV integration event locations using BEDTools (v. 2.30.0). Events and hotspots on chromosome X were excluded. The phase block borders of the HPV integration events were also identified by intersecting with BEDTools (v. 2.30.0), and the haplotype of the integration event was determined as the haplotype that contained the majority of reads at that integration event location. The *P* values of the DMR densities at HPV-associated DMR hotspots were produced by generating 1000 random genomic regions of the same size as the DMR hotspot, calculating the DMR density within the region for the test sample and all other samples, calculating the fold change of DMR density of the test sample vs. the mean of all other samples, and determining which percentile the test region’s fold change is to the 1000 random regions.

### Accession of gene expression data

The gene expression data for both the TCGA and HTMCP cohorts was downloaded from the GDC data portal on August 31st 2023 (https://gdc.cancer.gov/). The transcripts per million (TPM) normalized values were used to create an expression matrix of the 63 samples. No batch effects were observed between the expression of TCGA and HTMCP samples.

### Gene expression outliers and allele-specific expression analysis

The distance between protein coding genes (Ensembl 100 GRCh38) and integration event loci was calculated using BEDTools^54^ (v. 2.30.0). Genes that were within 1 Mb (+/−) of the integration events were tested to see if they were outliers, i.e. more than 1.5 IQR below Q1 or 1.5 IQR above Q3. The fold change of the outliers was calculated as log2(TPM of the tested sample/TPM of the median sample). ASE was determined using IMPALA^59^ (v. 1.0). Genes with ASE were defined as genes that had a major allele frequency threshold greater than 0.65 and an adjusted *P* value less than 0.05.

### Statistical analyses

No sample sizes other than the initial dataset were predetermined. Unless otherwise stated, all statistical tests correspond to two-sided tests. *P* value methods and multiple-test correction are reported in the text. Wilcoxon in the text refers to the Wilcoxon rank-sum test.

---

**Extended Data Figure 1:**
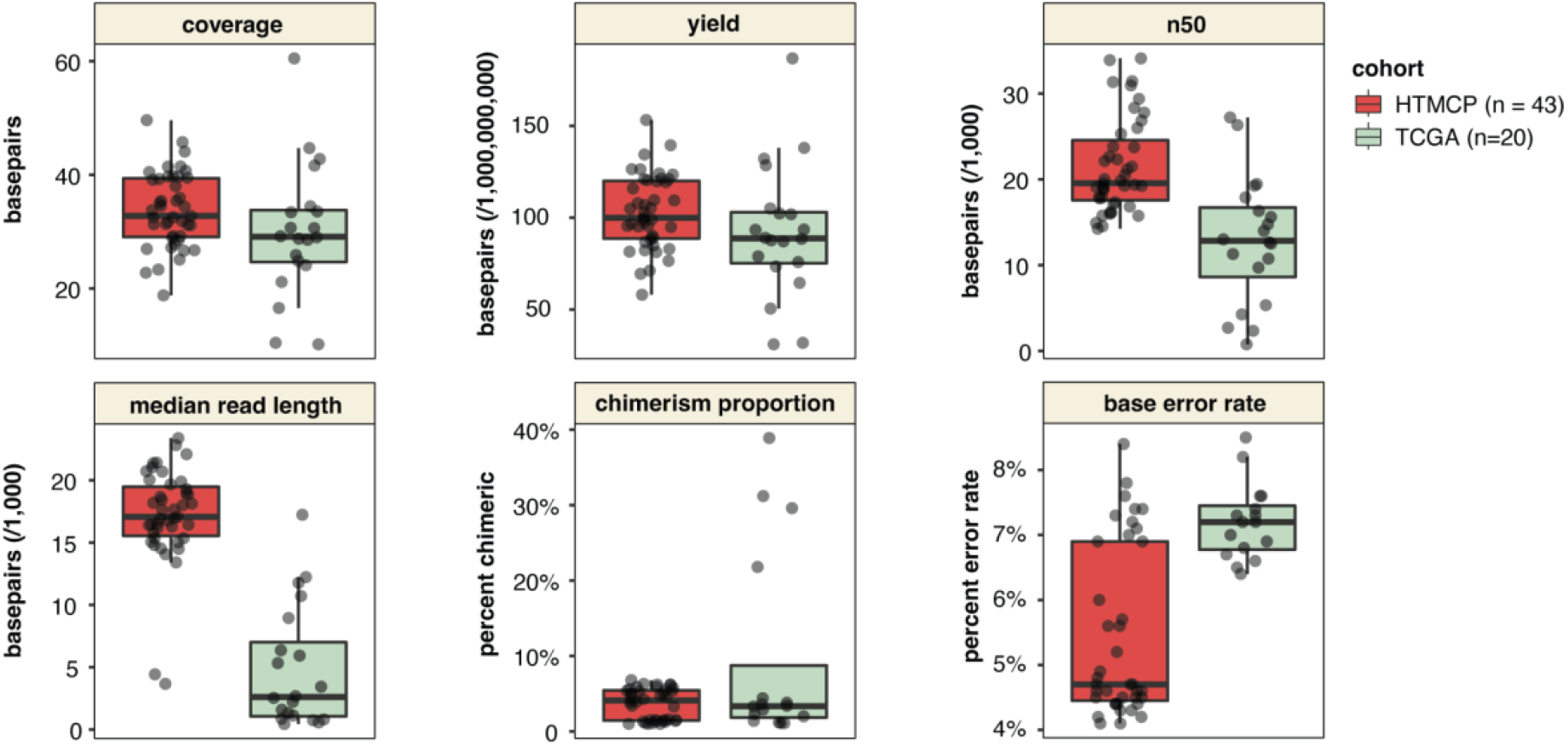
Sequencing statistics and quality control of ONT long-read sequencing. The coverage, yield, number of reads, n50, median read length, chimerism rate, and base error rate of the HTMCP and TCGA samples sequenced using whole genome ONT long-read technology. Box plots represent the median and upper and lower quartiles of the distribution; whiskers represent the limits of the distribution (1.5 IQR below Q1 or 1.5 IQR above Q3).

**Extended Data Figure 2:**
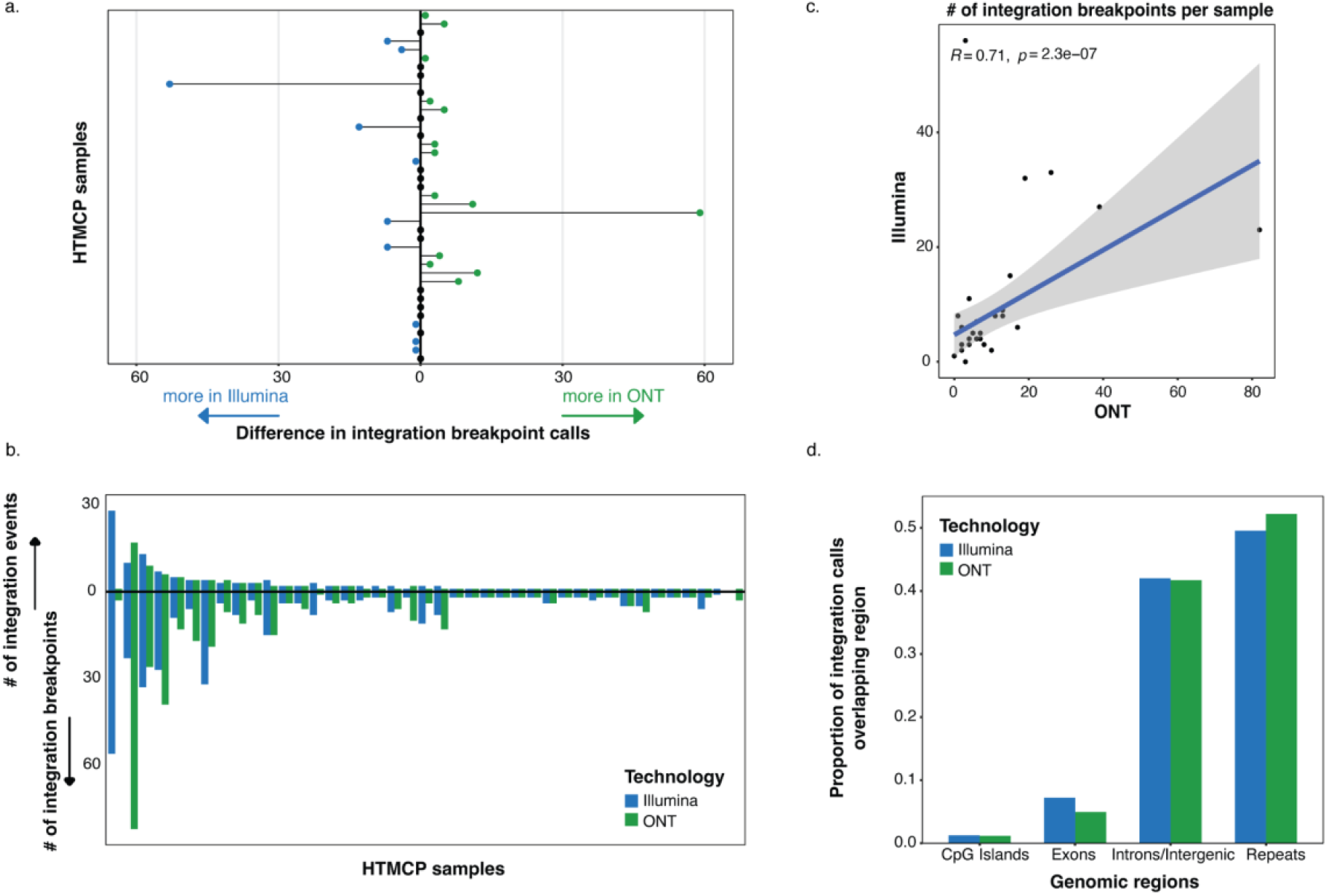
Comparison of HPV integration calling using Illumina short-read sequencing and ONT long-read sequencing. **(a)** The differences in the number of HPV integration breakpoint calls per sample in short-read (Illumina) and long-read (ONT) WGS. For samples where more breakpoints were detected by short-reads, the difference is shown to the left, and where more breakpoints were detected by long-reads, the difference is shown to the right. **(b)** The number of integration breakpoints and events in short-read (Illumina) and long-read (ONT) WGS. **(c)** The Spearman correlation between the number of HPV breakpoint calls per sample as determined by long-read sequencing (ONT) vs. short-read sequencing (Illumina). **(d)** Comparison of the genomic regions harboring breakpoint calls in the short-read (Illumina) and long-read (ONT) WGS.

**Extended Data Figure 3:**
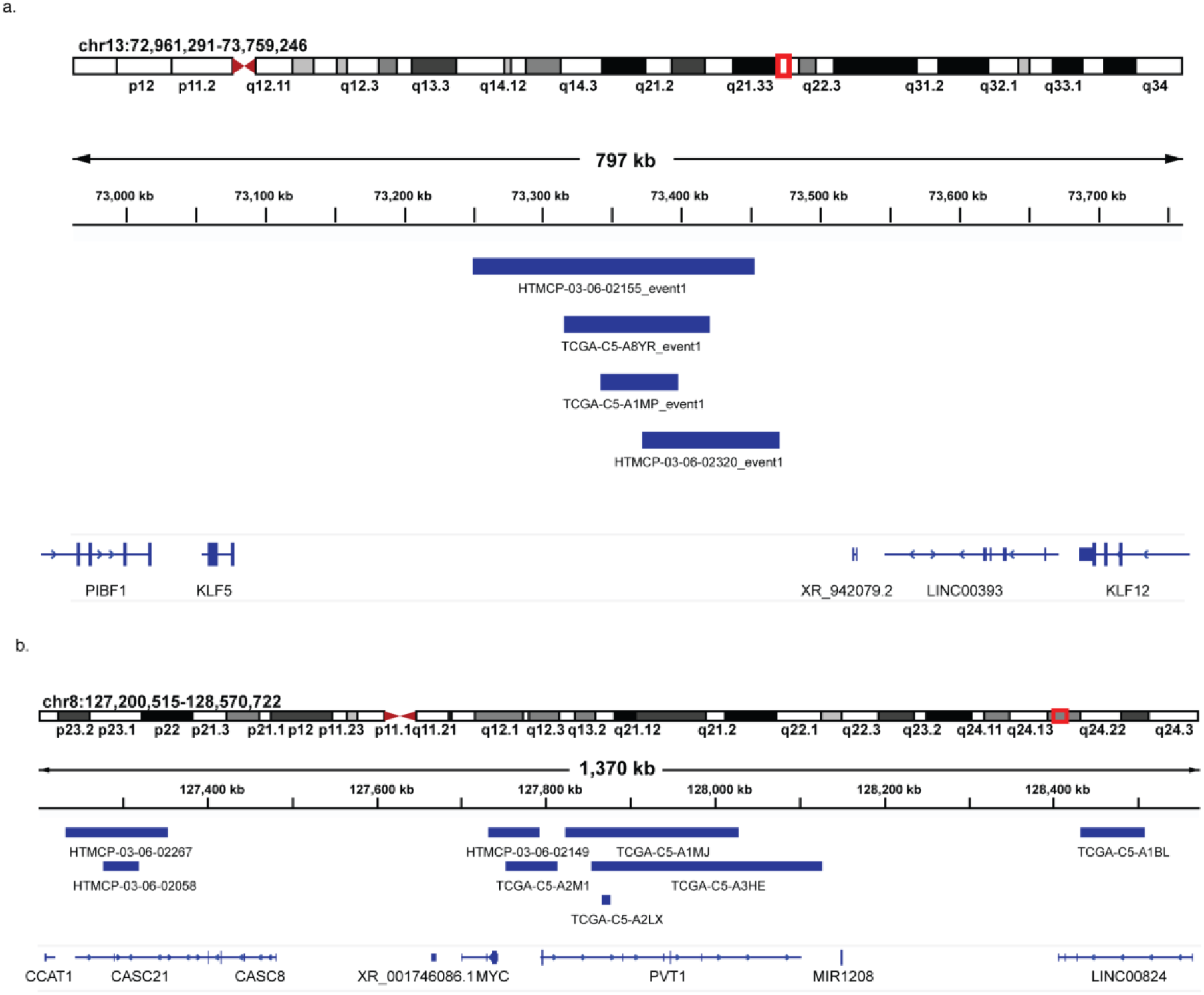
The distribution of events at loci recurrently affected by HPV integration. **(a)** The four overlapping eccDNA events in the intergenic region between *KLF5* and *KLF12* at the 13q22 locus. **(b)** The eight multi-breakpoint events in the 8q24 locus around *MYC*. The integration events were visualized in the Integrated Genome Browser, with the integration events represented as regions spanning the 5’ and 3’ integration breakpoints.

**Extended Data Figure 4:**
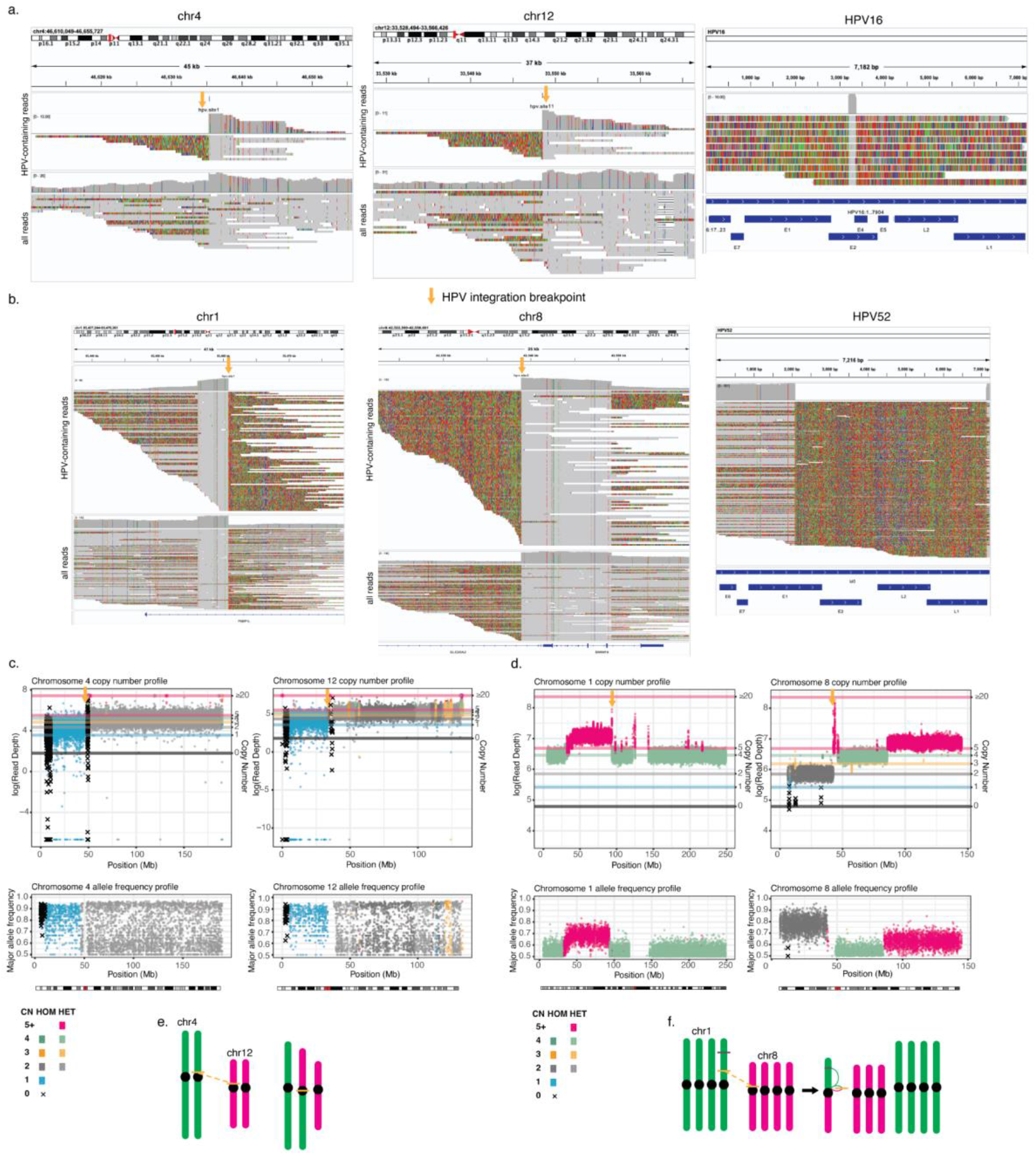
Visualization of translocation integration events. **(a,b)** Integrated Genome Browser snapshots with mismatched bases turned on and supplemental alignments included showing the HPV translocation events in (a) TCGA-A5-A1BL between chr4 and chr12, and in (b) HTMCP-03-06-02054 between chr1 and chr8. **(c,d)** The copy number profiles of (c) TCGA-A5-A1BL on chr4 and chr12, and (d) HTMCP-03-06-02054 on chr1 and chr8, as determined using Illumina WGS. **(e,f)** Schematic illustrating the inferred chromosome arm exchange resulting from the translocation integration events in (a) and (b). Orange arrows and lines denote the HPV integration breakpoints.

**Extended Data Figure 5:**
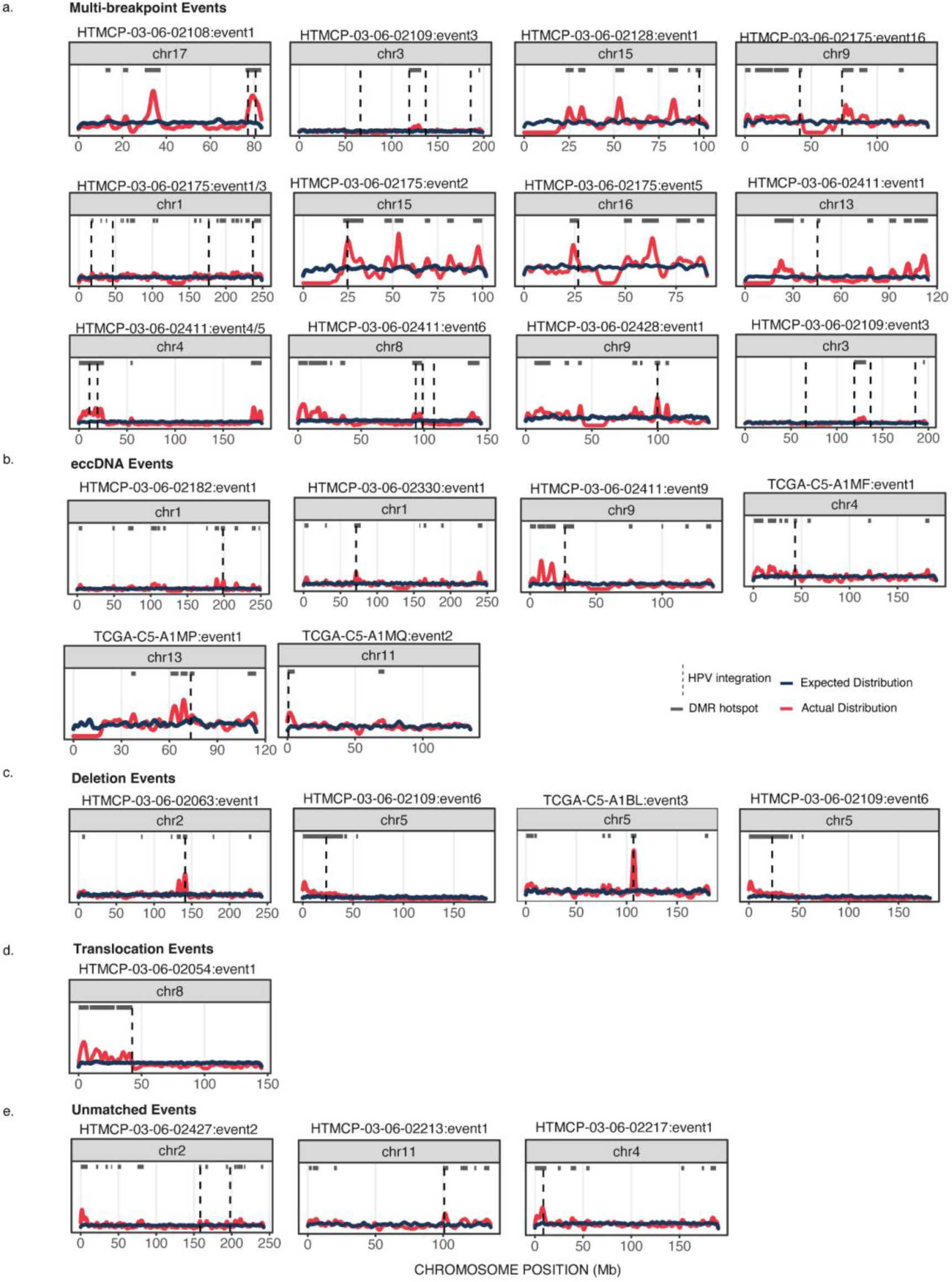
Additional examples of DMR density hotspots on chromosomes harboring HPV integration. **(a-e)** Chromosome-wide DMR density plots of all HPV integration events overlapping DMR hotspots, including (a) multi-breakpoint events, (b) eccDNA events, (c) deletion events, (e) translocation events, and (e) events with a single unmatched HPV breakpoint, i.e. undetermined integration category. The dotted lines indicate sites of HPV integration.

**Extended Data Figure 6:**
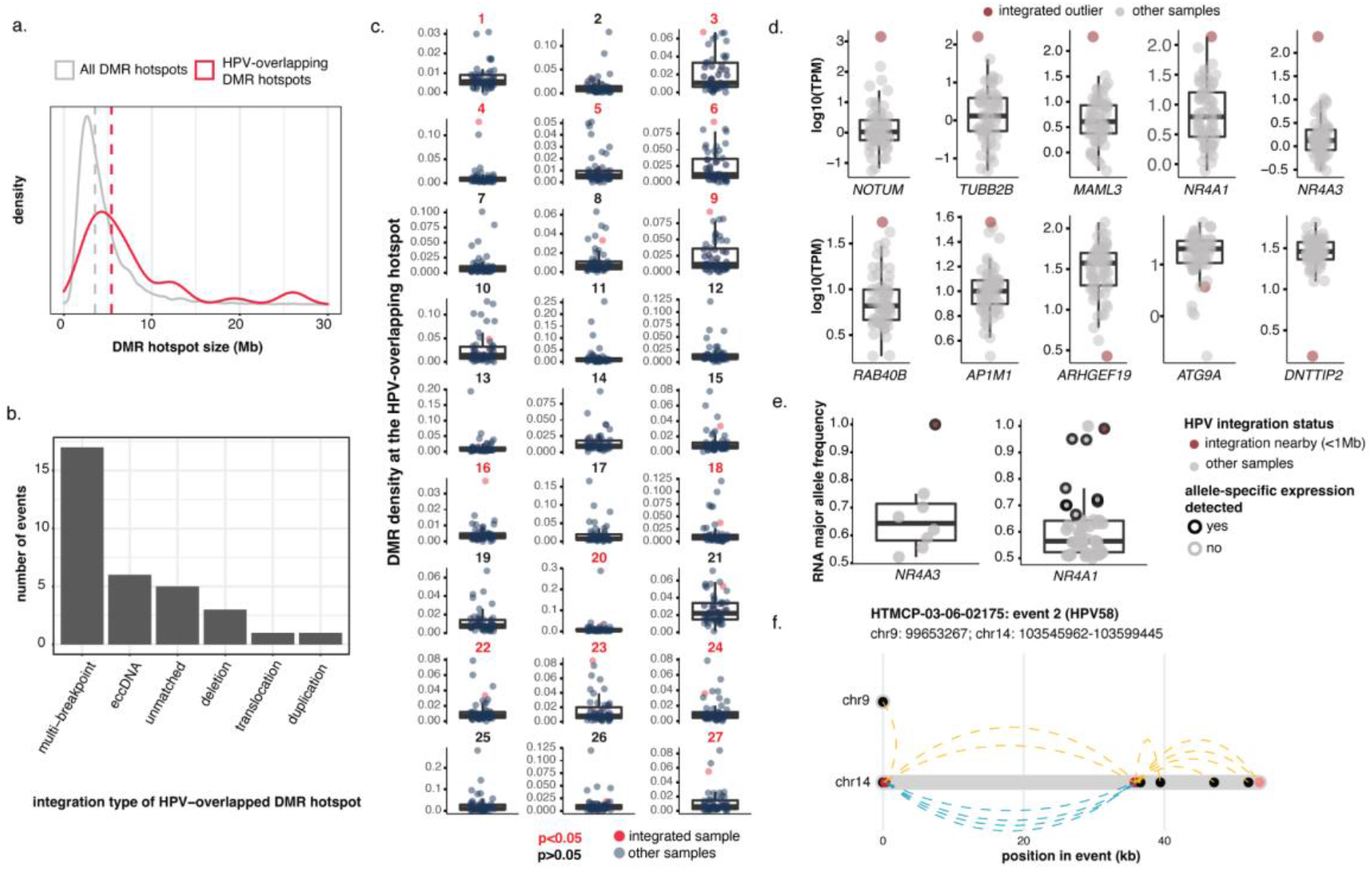
Additional information on the DMR hotspots and upregulated genes around HPV integration sites. **(a)** The genomic span of HPV-associated DMR hotspots compared to all DMR hotspots across the cohort. The dotted lines denote the median of each distribution. **(b)** The frequencies of HPV integration events overlapping a DMR hotspot across the different integration categories. **(c)** Statistical testing of the 27 integration events overlapping DMR hotspots. For each integration event, the DMR density in the integrated sample was compared to all other samples in the cohort. *P* values were determined by a permutation test across the genome with the window size equal to the size of the DMR hotspot. **(d)** Gene expression differences between integrated samples with outlier gene expression nearby (+/− 1Mb) HPV integration and other samples without HPV integration in that region. **(e)** The RNA-seq major allele frequency and ASE of *NR4A1* and *NR4A3* in the sample with HPV integration nearby compared to the other samples. **(f)** The breakpoint structure of HTMCP-03-06-02428’s event 1 neighboring *NR4A3* on chromosome 9. Box plots represent the median and upper and lower quartiles of the distribution; whiskers represent the limits of the distribution (1.5 IQR below Q1 or 1.5 IQR above Q3).

**Extended Data Figure 7:**
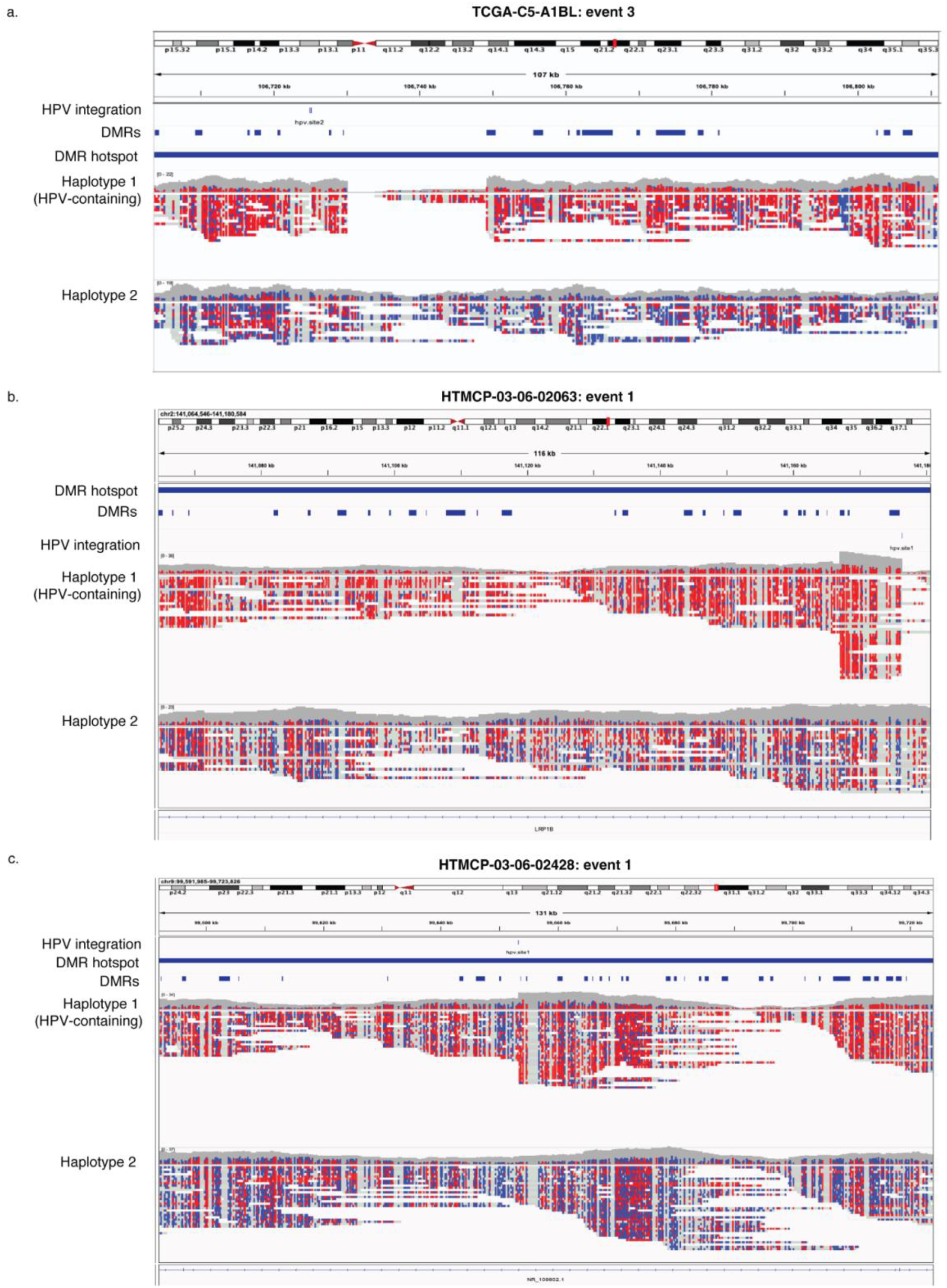
Examples of differential methylation at HPV integration events overlapping DMR hotspots. **(a-c)** Integrated Genome Browser snapshots of the genomic regions surrounding HPV integration events, including two deletion events, (a) TCGA-A5-A1BL: event 3 and (b) HTMCP-03-06-02063: event 1, and one multi-breakpoint event, (c) HTMCP-03-06-02428: event 1. The reads are separated by haplotype and CpGs are colored by methylation status (red = methylated, blue = unmethylated). Tracks showing the HPV integration breakpoints, the DMRs, and the DMR hotspot regions are also included in the snapshots.

**Extended Data Figure 8:**
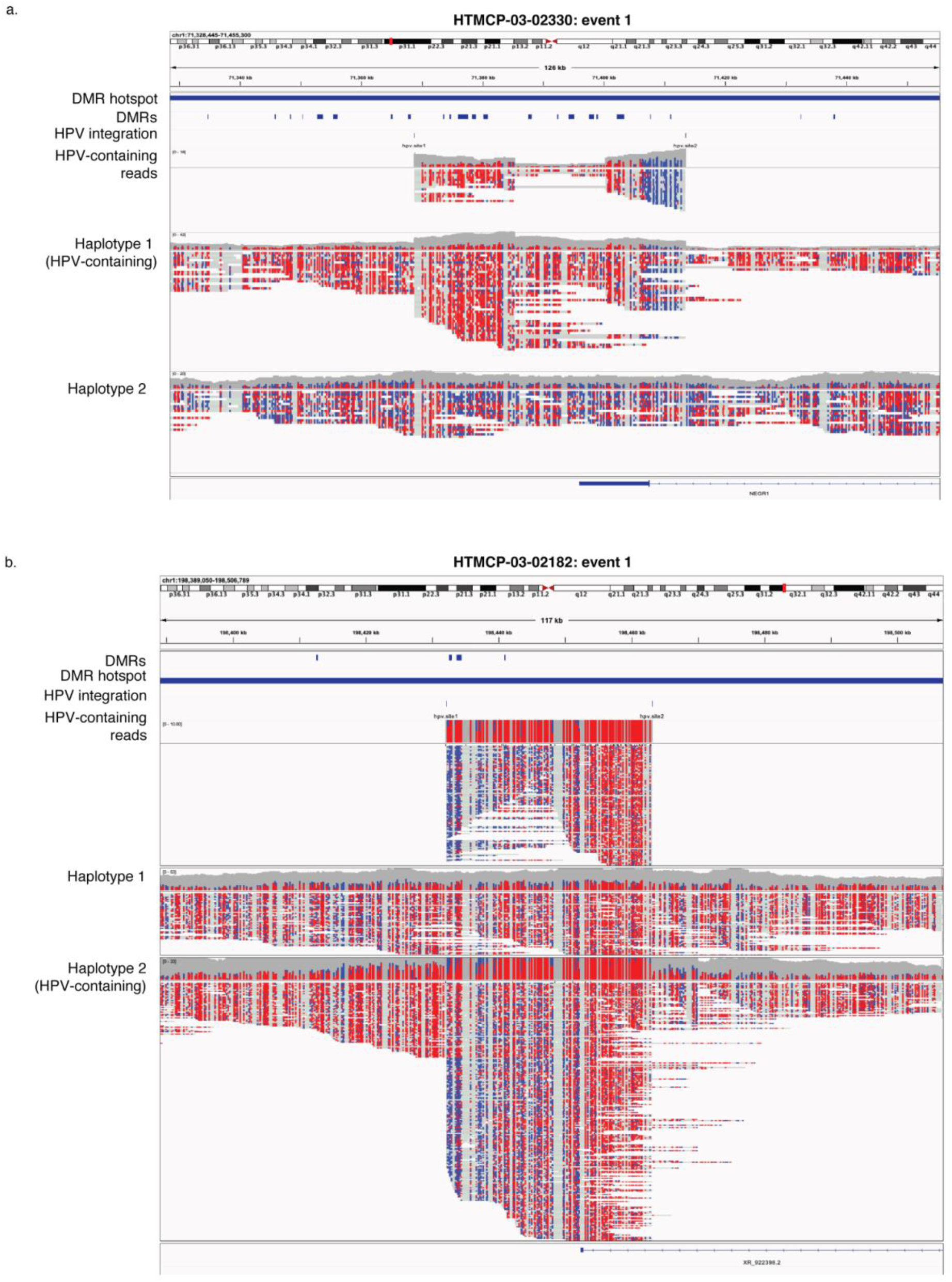
Examples of differential methylation at eccDNA events overlapping DMR hotspots. **(a,b)** Integrated Genome Browser snapshots of genomic regions surrounding eccDNA integration events from (a) HTMCP-03-02330: event 1, and (b) HTMCP-03-02182: event 1. The reads are separated by haplotype and CpGs are colored by methylation status (red = methylated, blue = unmethylated). Tracks showing the HPV integration breakpoints, the methylation DMRs, and the DMR hotspot regions are also included in the snapshots.

